# Complex networks of miRNA-transcription factors mediate gene dosage compensation in aneuploid cancer

**DOI:** 10.1101/2020.01.31.928507

**Authors:** ManSai Acón, Guillermo Oviedo, Edwin Baéz, Gloriana Vásquez-Vargas, José Guevara-Coto, Andrés Segura-Castillo, Francisco Siles-Canales, Steve Quirós-Barrantes, Pedro Mendes, Rodrigo Mora-Rodríguez

## Abstract

Cancer complexity is consequence of enormous genomic instability leading to aneuploidy, a hallmark of most cancers. We hypothesize that dosage compensation of critical genes could arise from systems-level properties of complex networks of microRNAs (miRNA) and transcription factors (TF) as a way for cancer cells to withstand the negative effects of aneuploidy. We studied gene dosage compensation at the transcriptional level on data of the NCI-60 cancer cell line panel with the aid of computational models to identify candidate genes with low tolerance to variation in gene expression despite high variation in copy numbers. We identified a network of TF and miRNAs validated interactions with those genes to construct a mathematical model where the property of dosage compensation emerged for MYC and STAT3. Compensation was mediated by feedback and feed-forward motifs with 4 miRNAs and was dependent on the kinetic parameters of these TF-miRNA interactions, indicating that network analysis was not enough to identify this emergent property. The inhibition of miRNAs compensating MYC suggest a therapeutic potential of targeting gene dosage compensation against aneuploid cancer.

## Introduction

Cancer is a heterogeneous group of diseases characterized by out of control cell growth and division sharing several hallmarks, including 8 functional capabilities and 2 enabling characteristics. Among those enabling characteristics, genome instability is one of the drivers of cancer development (Hanahan & Weinberg, 2011). Indeed, cancer robustness is enabled at the tumor cell population level by heterogeneity in therapy responses, which is driven by genomic instability (Kitano, 2004), specially by aneuploidy: gains and losses of whole or partial chromosomes.

Genomic instability leads to aneuploidy in most cancers: 90% of solid tumors and 75% of hematologic tumors present aneuploidy, which was considered by early reports as the driving force of cancer genomic evolution (Sheltzer & Amon, 2011). Indeed, a report indicated that aneuploidy is the major source of autocatalytic genomic instability, where the cells with the highest aneuploidy have also the highest instability of their genomes (Duesberg, Rausch, Rasnick, & Hehlmann, 1998). Aneuploidy seems to have a role in oncogenesis because: i) it appears before malignant transformation, ii) there are clonal genetic alterations that compromise the fidelity of chromosomal segregation and iii) there are inherited mutations in checkpoints leading to aneuploidy and predisposing to cancer (Sheltzer & Amon, 2011).

Nevertheless, aneuploidy is lethal for normal cells and entire organisms, representing the most frequent cause of abortions and mental retardation in humans. All monosomies (loss of one chromosome) and 20 out of 23 possible trisomies (gain of one additional chromosome) are lethal and out of the other three, only trisomy 21 can survive adulthood (Down Syndrome). This means that aneuploidy is a barrier to a successful development of the embryo and leads to many defects at the cellular level. Aneuploidy decreases cell proliferation and viability, increases proteotoxic stress, metabolic requirements, lactate production and induces recombination defects leading to genomic instability (Sheltzer & Amon, 2011). The most likely explanation for all those negative effects of aneuploidy is given by the alteration of the gene dosage: gains or losses of entire chromosomes immediately alter the doses of hundreds of genes in the cell, leading to unbalanced load of critical proteins, altering energetic requirements and protein homeostasis (Sheltzer & Amon, 2011).

Although lethal to normal cells, aneuploidy is a hallmark of most advanced cancer cells. Therefore those cells must have developed mechanisms to minimize the negative effects of aneuploidy. First, aneuploidy autocatalizes genomic instability of transforming cancer cells leading to many unstable karyotypes and cell death due to an error catastrophe (Solé & Deisboeck, 2004). However, in rare occasions, a specific combination of various alterations is met that overcomes those error thresholds in cancer evolution, leading to malignant cells that are able to survive aneuploidy and genomic instability. This bottleneck in cancer evolution represents a gate to evolve malignant karyotypes leading to drug-resistance and metastasis (L. Li et al., 2009). However, this evolution is not unrestricted. Within the chaos of chromosomal instability, some conserved patterns in the karyotypic configurations suggest the presence of a stable mechanism, which function has to be maintained to ensure survival: i) there are specific aneusomies at different stages of cell transformation (Fabarius, Li, Yerganian, Hehlmann, & Duesberg, 2008), ii) there are clonal karyotypes evolving during the cell passages (Fabarius, Willer, Yerganian, Hehlmann, & Duesberg, 2002), iii) the cancer causing karyotypes have a chromosomal equilibria between destabilizing aneuploidy and the stabilizing selection for oncogenic function (L. Li et al., 2009), and iv) a large scale study revealed 2 distinct pathways to aneuploidy where the cells gain or lose chromosomes to restore the balance of their altered proteins and maintain viability (Ozery-Flato, Linhart, Trakhtenbrot, Izraeli, & Shamir, 2011). Therefore, despite the genomic instability of cancer, these observations suggest the existence of a hidden karyotypic pattern responsible for maintaining a stable mechanism to cope with the negative effects of aneuploidy. Finding these karyotypic configurations from genomic observations represents a huge challenge of pattern recognition.

It is therefore unknown how cancer cells deal with so much aneuploidy whereas normal cells are very sensitive. A possible explanation is given by the hypothesis of gene dosage compensation, a mechanism that has been described very early for other organisms to compensate the negative effects of aneuploidy (Devlin, Holm, & Grigliatti, 1982). Indeed, the concept of gene dosage compensation, or gene dosage balance, is a widespread phenomenon that was discovered in the early days of genetics and there is accumulating evidence that it has an effect on gene expression, quantitative traits, aneuploid syndromes, population dynamics of copy number variants and differential evolutionary fate of genes after partial or whole-genome duplications (for review see (Birchler & Veitia, 2012)). The effects of gene dosage compensation are hypothesized to result from stoichiometric differences among members of macromolecular complexes, the interactome, and signaling pathways (Birchler & Veitia, 2012; Veitia, Bottani, & Birchler, 2008). Gene dosage compensation represents a compensatory mechanism that may ameliorate the imbalanced gene expression and restore protein homeostasis in aneuploid cells. The multiple consequences of aneuploidy are mainly regulated at the protein level due to a side-effect of protein folding defects and increased protein degradation by proteosome and autophagy (Donnelly & Storchová, 2014). An approach to identify genes with dosage compensation by increasing the copy number of individual genes using the genetic tug-of-war technique showed that approximately 10% of the genome shows gene dosage compensation, and consists predominantly of subunits of multi-protein complexes, which are regulated in a stoichiometry-dependent manner (Ishikawa, Makanae, Iwasaki, Ingolia, & Moriya, 2017), although this approach was designed to identify compensated genes at the protein level only.

In aneuploid cancer it has been shown that messenger RNA (mRNA) levels generally correlate well with an increased DNA copy number (gene dosage) but these changes are not reflected at the protein level for several genes (Stingele et al., 2012). Indeed, gene dosage compensation in cancer cells has been recently (Brennan et al., 2019) demonstrated at the protein level, whereby protein aggregation mediates the stoichiometry of protein complexes in aneuploid cells. In that study they show that excess subunits are either degraded or aggregated and that protein aggregation is nearly as effective as protein degradation at lowering the level of functional proteins (Brennan et al., 2019).

The regulation of gene transcription might be another effective mechanism to compensate the gene dosage changes in aneuploid cells, as it maintains the stoichiometry and preserves the energy required for transcription, translation, and eventual degradation of the extra proteins. However, most of the transcriptional analyses of model aneuploid cells in budding and fission yeasts, plants, mice and human cells have indicated that mRNA levels scale with gene copy number, excepta few Drosophila genes that appear to be the vestige of a sex-determining silencing mechanism (Donnelly & Storchová, 2014). Actually, the inactivation of the X chromosome in female mammals represents an early reported mechanism of gene dosage compensation mediated by the Xist non-coding RNA (Park & Kuroda, 2001). Nevertheless, a previous report of the insertion of an additional chromosome 5 revealed that most proteins coded on the extra chromosomes are more abundant than proteins from diploid chromosomes indicating that there is no general efficient mechanism for ‘gene dosage compensation’ in this system. However, some specific proteins are maintained at diploid levels, specially those corresponding to kinases and ribosomal subunits. Most of these genes are compensated at the protein level but a few others are also compensated at the mRNA level (Stingele et al., 2012). In addition, a report on aneuploid wild yeast isolates showed gene-dosage compensation in 10-30% of amplified genes compared to isogenic or closely related euploid strains and that aneuploidy did not lead to growth defects. In fact, the authors propose that gene dosage compensation is most likely due to feedback control and enables a rapid karyotypic evolution in yeast. They also predicted that dosage compensation occurs at genes that are most toxic when overexpressed and that their expression may also be under greater evolutionary constraint (Hose et al., 2015).

Feedback control that buffers the mRNA levels of amplified or deleted chromosomal regions has been already suggested in naturally occurring yeast strains (Kvitek, Will, & Gasch, 2008) as well as in lager brewing yeast (Bond, Neal, Donnelly, & James, 2004). Moreover, it has been shown that a mechanism based on an incoherent feedforward motif enables adaptive gene expression in mammalian cells. Using synthetic transcriptional and post-transcriptional incoherent loops, the authors found that the gene product adapts to changes in DNA template abundance, supporting a previously hypothesized endogenous role in gene dosage compensation for such motifs (Bleris et al., 2011; Shimoga, White, Li, Sontag, & Bleris, 2014). In endogenous transcription networks, the interactions of miRNAs and transcription factors have been reported to assemble those kind of complex motifs including negative feedback loops, positive feedback loops, coherent feed-forward loops, incoherent feed-forward loops, miRNA clusters and target hubs leading to non-linear, systems-level properties such as bistability, ultrasensitivity and oscillations (Lai et al., 2013; Vera, Lai, Schmitz, & Wolkenhauer, 2013).

miRNAs are small endogenous RNA molecules that bind mRNAs and repress gene expression (Fabian, Sonenberg, & Filipowicz, 2010). miRNAs are one of the most predominantly represented non-coding RNA (ncRNA) groups in clinical research. A typical miRNA is processed from a long primary RNA sequence to a short mature functional transcript around 22 nucleotides in length. A common characteristic of a miRNA is its ability to pleiotropically target potentially hundreds or even thousands of genes (Hanna, Hossain, & Kocerha, 2019) and their target genes can also be regulated by several miRNAs (Ritchie, Rasko, & Flamant, 2013). Indeed, current estimates point to the human genome containing 1917 annotated hairpin precursors, and 2654 mature sequences of miRNAs (Kozomara, Birgaoanu, & Griffiths-Jones, 2019), estimated to directly regulate >60% of human mRNAs (Kim et al., 2016). In consequence, there is a good possibility that miRNA-transcription factor interactions may regulate the expression of genes amplified or deleted in cancer. Therefore, we hypothesize that gene dosage compensation in cancer can be mediated, at least in part, by the emerging properties of complex miRNA-TF networks, controlling the expression of genes that have altered copy number.

In the present work we set out to investigate whether we could identify such a mechanism by analyzing genomic and transcriptomic data of the NCI60 cell panel. We have indeed identified a cluster of genes with low tolerance to variation in expression level despite having a high variation in copy numbers across the cell lines. These genes are distributed along several chromosomal locations and indeed, their copy numbers have positive or negative correlation coefficients with the expression levels of several miRNAs and transcription factors. Most of these correlations do not correspond to direct target interactions suggesting that dosage compensation could be mediated by a complex network of miRNAs and transcription factor interactions. Due to the complexity of the underlying biochemical network, identification of such mechanisms and their molecular components is not trivial. We developed a computational platform for that purpose, including bioinformatics and dynamical modeling tools. This platform starts with various genomic data and, using existing knowledge, builds a network of miRNA and TF interactions, embodied as a mathematical model of ordinary differential equations (ODE) using mass action kinetics. The parameters of this model are estimated by fitting to the NCI60 data, and we were able to reproduce dosage compensation for MYC and STAT3, and identify several candidate miRNAs and TF that likely mediate this phenomenon. Using the ODE model for steady state simulations and perturbation experiments, we were able to reduce the model complexity and identify a minimal model of gene dosage compensation for MYC and STAT3 involving 4 miRNAs with redundancy in their function of dosage compensation.We validated this model by inhibition of those miRNAs in an experimental model of colon cancer, which led to increasing levels of cytotoxicity as MYC copy number increases. Thus, the disruption of the mechanism of gene dosage compensation leads to cell death. This dosage compensation mechanism for MYC and STAT3 may represent a sentinel core for the dosage compensation of other genes with copy number alterations happening simultaneously with those of MYC and STAT3. The kinetic parameters of these TF-miRNA interactions determine the capability of these network motifs to achieve gene dosage compensation. Searching for similar motifs among the putative interactions reported for other candidate target genes, we could suggest a putative mechanism for the dosage compensation of STAT5B and FOXC1 as well.

We propose the existence of a regulatory network mediated by miRNAs that compensates for gene dosage changes in aneuploid cancer cells. We suggest that the manipulation of specific nodes of this miRNA-based regulatory network could block gene dosage compensation, representing a specific target against aneuploid cancer.

## Results

### Candidate genes under transcriptional gene dosage compensation are present across the cancer genomes of the NCI-60 panel

To identify genes under possible dosage compensation, we compared copy number, gene expression and proteomic data of all genes in the NCI60 panel. We considered input data including high resolution copy Number Variation data (DNA) of the NCI-60 Cancer Cell lines from 4 different platforms (Bussey et al., 2006), the Gene Transcript (RNA) Average Intensities of 5 Platforms (Gmeiner, Reinhold, & Pommier, 2010), and the protein levels (Protein) of a global proteome analysis of the NCI-60 cell line panel (Gholami et al., 2013). Figure 1A-left shows the variation of the absolute values of DNA copy number, RNA expression and Protein expression. Next, we calculated the average RNA or protein expression of the diploid cell lines (copy number between −0.25 and 0.25) for each gene to obtain the average diploid expression for every gene. Afterwards, we normalized all expression data against this diploid average expression for each gene and calculated log2 transformed values. Using this approach, we could directly and simultaneously compare the variation in DNA Copy Number variation, RNA and Protein expression across the NCI60 panel to look for genes with high variations in DNA copy numbers but a low tolerance to variation in RNA or protein expression as they could represent candidate targets under dosage compensation (Figure 1A). Indeed, we plotted the Standard Deviation (SD) of the DNA, RNA and Protein values across the 59 cell lines and observed that several dots separated from the main gene population as they present high SD of DNA levels (Figure 1B).

**Figure 1.**
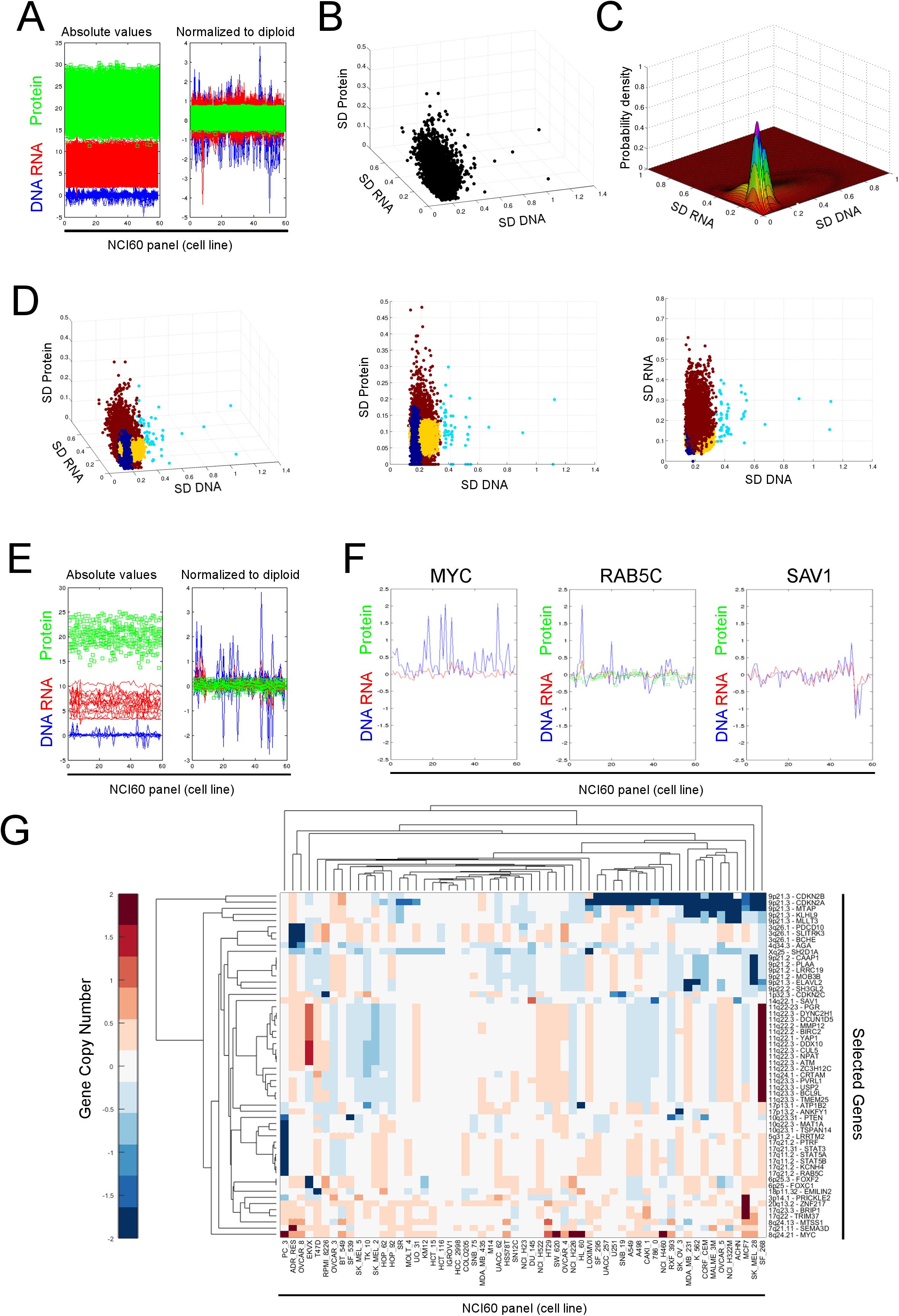
Identification of candidate genes under dosage compensation. A. Input data of gene copy number (DNA), gene expression (RNA) and protein levels (protein) of the NCI60 panel. The absolute values are shown on the left panel and the right panel corresponds to the log2 values normalized to the averaged RNA and protein of the diploid cell lines for the respective gene (Normalized to diploid). B. Standard deviations (SD) of the DNA, RNA and Protein levels for each gene across the 59 cell lines of the NCI60 panel. C. Gaussian Mixture Model to identify a cluster of subpopulation of genes with high SD DNA and low SD RNA and/or low SD Protein (white arrow). D. Gene Clustering according to the model in C (left). The cyan cluster contains candidate genes under dosage compensation, characterized by high SD DNA, low SD Protein (middle) and low SD RNA (right). E. Absolute and Normalized values of selected candidate genes under dosage compensation. F. Examples of candidate genes under dosage compensation (MYC and RAB5C) compared to a non-candidate gene (SAV1). G. Clusters of candidate genes according to their copy number variations across the 59 cell lines of the NCI60 panel.

To obtain a good separation of candidate compensated genes from the main cluster, we implemented a Gaussian mixture model (GMM) with these data (Figure 1C). After classification, we could observe one group of genes with high SD of DNA copies and relatively low SD of RNA and/or protein expression values (Figure 1D). This cluster showed the expected behavior with high amplitudes in DNA values but very low variation in RNA or protein expression (Figure 1E). For example, MYC presents high frequency of amplifications in the NCI60 panel without the corresponding increase in RNA levels (Figure 1F right). A similar behavior is observed for RAB5C for both RNA and protein levels despite its copy number variations (Figure 1F middle), compared to a gene such as SAV1 where copy number and RNA expression are well correlated (Figure 1F right). Next, we discarded those genes with orthologues in X/Y chromosomes since they cannot be differentiated using microarray techniques, and obtained a list of 56 gene candidates with putative dosage compensation at the transcript level. In some cases there was also protein data confirming this behavior but there was no candidate with putative compensation at the protein level only.

We next aimed to correlate the behavior of the copy number variations among these candidate genes to their corresponding chromosomal locations across the NCI60 panel. The clustergram in Figure 1G shows different clusters suggesting a similar behavior of copy number variations. As expected, many of these genes are correlated by their genomic locations, especially gene clusters belonging to the chromosomal bands 3q26, 6p25, 8q24, 9p21-22, 10q22-23, 11q22-24, 17q11 and 17q21-23. These data suggests the existence of genes with low tolerance to variation in their RNA expression despite high copy number variation across the NCI60 panel, partially related by common chromosomal locations.

We hypothesized, at this point, that the inhibition of a dosage compensation mechanism would release the brake imposed on the expression of all these additional copies leading to significant over-expression of these genes. If the dosage compensation of these genes was favored during cancer evolution, its over-expression could potentially lead to death of the cell lines with these specific amplifications. Therefore, we focused our next steps on the description of possible mechanisms involved in dosage compensation of amplified genes. For example, MYC presents 27 compensated amplifications with very similar expression levels even with 8 genetic copies. Therefore, we reduced our list of candidates by discarding genes with high DNA variation due mostly to deletion (more than 6 cell lines with compensated deletions) and kept only genes with at least 6 cell lines with compensated amplifications. This approach reduced our list to 21 candidate genes with dosage amplifications (Expanded View Figure 1) potentially compensated at the transcript level across the NCI60 panel.

### A mathematical model of gene dosage compensation mediated by a network of miRNA-transcription factor interactions

We asked whether dosage compensation mechanism could be mediated by systems-level properties arising from a complex regulatory network of gene expression. Since there is expression data available of miRNAs (Blower et al., 2007) and transcription factors (TF) (Gmeiner et al., 2010) for the NCI60 panel, we performed a correlation analysis of the copy numbers of target genes with the expression levels of miRNAs and transcription factors in order to identify possible regulators responding to the dosage of our candidate genes (Expanded View Figure 2). There are both miRNAs and TF with positive or negative correlation to the candidate copy numbers (Expanded View Figure 2A), although most of those correlations do not correspond to the direct reported interactions (Expanded View Figure 2B). Using all those interactions (see Expanded View Section) we constructed a network of putative and validated regulatory interactions connecting all candidate genes (Expanded View Figure 2C). However, from the construction of this network topology is easy to infer that this network could sense changes in copy number of target genes only if they also have a TF-function (red colored interactions, Expanded View figure 2C). Only a gene candidate with TF function will trigger a signal that can be propagated throughout the network and back if this gene has a regulatory loop. This loop would compensate gene expression in response to changes in gene dosage.

The regulatory motifs with potential systems-level properties to mediate dosage compensation are widely present within this putative network (Expanded View section). Nevertheless, a gene-dosage response mediated by one or several of these regulatory loops will depend on the strength of the interactions (i.e. their kinetic parameter values). Due to the high complexity of this miRNA/TF regulatory network, we envisaged the construction of a large scale mathematical model in order to gain insight into a possible mechanism of gene dosage compensation mediated by any of the many regulatory loops identified in this network. A quantitative systems-level approach for the calculation of the goodness of fit of a mathematical model describing the interactions enabled us to assess the feasibility of a proposed network topology for gene dosage compensation.

First, we reduced the network complexity to include only the target genes with TF-function as they are the only genes for which a copy number variation can be sensed. This resulted in a network of 517 nodes and 44016 arcs of putative and validated interactions. Since we collected about 65.000 experimentally validated interactions out of HTRI, Pazar, Transmir and Mirtarbase, we decided to include only experimentally validated interactions to further reduce model complexity. This led to a network of 78 nodes and 578 arcs of experimentally validated interactions. After this simplification, several target genes became dead ends, since no validated interactions are reported influencing other nodes of the network. Therefore, we removed them, further reducing the model to include only 4 target genes with TF-function and validated interactions influencing other nodes of the network: MYC, STAT3, STAT5A and STAT5B. This led to a network of 65 nodes and 506 arcs. The nodes include 45 TFs and 20 miRNAs. The arcs formed 16 feedback loops, 28 coherent feed-forward loops and 45 incoherent feed-forward loops (Figure 2A).

**Figure 2.**
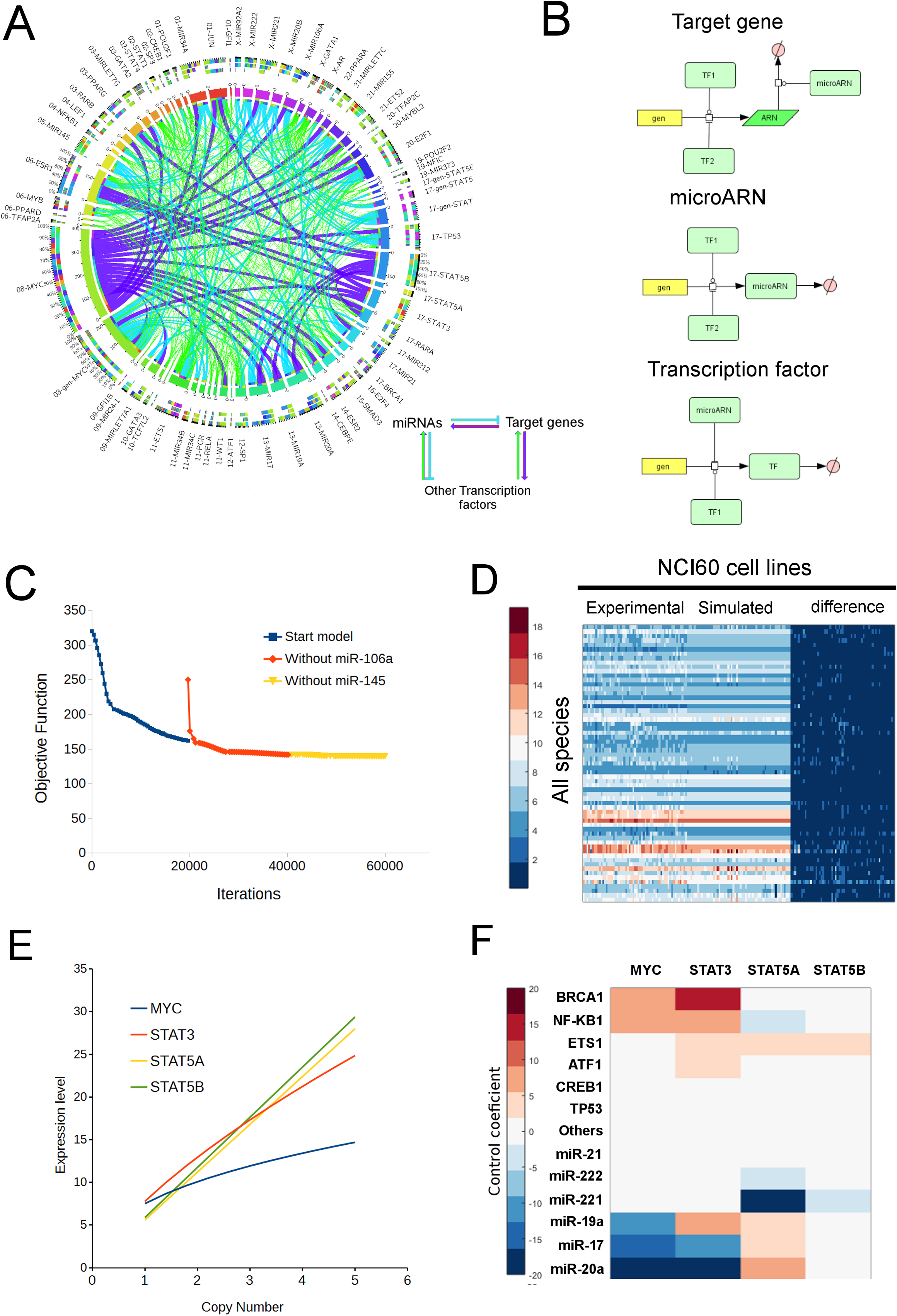
A mathematical model driven by NCI-60 data leads to a phenotype of gene dosage compensation for MYC and STAT3. A. Network of miRNA-TF interactions for MYC, STAT3, STAT5A and STAT5B. B. Schematic representations of the modeled interactions for the target gene, miRNAs and associated TFs. C. The results of the parameter estimation task to fit the ODE mathematical model to the experimental data upon different modifications led to a reduction in the objective function (difference between modeled and experimental data). D. The graphical comparison of modeled and experimental data shows no significant differences. E. Results of parameter scan on the values of copy number for the candidate target genes showing gene dosage compensation for MYC and STAT3. F. The sensitivity analysis points to some candidate regulatory molecules controlling the concentrations of MYC and STAT3.

Second, we built an automated platform to generate ODE mathematical models for COPASI based on our candidate network topology. To do so, we introduced ODE expressions for the schematic representations of the modeled interactions for the target gene, miRNAs and associated TFs (Figure 2B) and translated them into differential equations. The biochemical model includes 65 species, 130 reactions and 305 parameters. The mathematical description of these reactions was represented using mass action kinetics under the assumption that each cell line of the NCI-60 panel represents a different steady state of this model depending only on the copy numbers of the genes in the network. The model parameters were estimated to fit the model to the data using the Parameter Estimation function of COPASI with the Hooke-Jeeves algorithm of optimization (Hooke & Jeeves, 1961). We increased the weight of the objective function of the expression of the 4 target genes in order to favor the search of model parameter values leading to dosage compensation. Model refinement was performed by interrupting the fitting process after several days without any significant progress. At these periods of time, we calculated the goodness-of-fit for each species and performed a metabolic control analysis. We calculated the p-values (t-Student to compare simulated with experimental data) for the goodness-of-fit for the first resulting model, which presents a good fitting for most species except for miR106a (p = 0.012). After removing miR106a, the resulting model achieved a lower objective function value (better fit) but miR145 had the worse goodness-of-fit (p=0.075) without significant control on the target genes. We decided to remove also miR145 from the model and continued the parameter estimation to obtain an optimized model with a lower objective function (figure 2C). For this last fit, there was no statistical difference of the values of the simulated species compared to the experimental data (p>0.1, figure 2D), obtaining thereby a mathematical model describing the experimental data of our target candidate and their associated miRNAs and TFs.

Furthermore, we asked whether the resulting model is able to compensate the gene dosage of our candidate target genes in response to an increase in gene copy numbers. In order to ascertain this, we performed a parameter scan using COPASI for the copy number of the 4 target genes individually. As shown in figure 2E, increasing the copy number of MYC from 1 to 5 leads only to an increase of 1.95 fold in MYC expression. For STAT3, an increase from 1 to 5 in copy number leads to an increase of 3.19 fold in STAT3 expression. This was not the case for STAT5A and STAT5B, which copy number increases were not compensated in the model. We next performed a metabolic control analysis using COPASI to identify candidate miRNAs or TFs with high positive or negative regulatory control on the expression level of the 4 target genes. We plotted the 6 miRNAs with the highest negative control and the 6 TFs with the highest positive control (figure 2F). The results showed that MYC levels are mainly controlled by an interplay between miR-17, miR20a, miR19a and BRCA1 and NF-KB1. STAT3 levels are mainly controlled by miR-17, miR-20a, miR19a and BRCA1, NF-KB1, ETS1 and ATF1. Although STAT5A and STAT5B were not compensated upon changes in their copy numbers, they also present control in their expression levels by some of the same miRNAs and TFs, suggesting that the mechanism of MYC and STAT3 compensation could also impact the levels of STAT5A and STAT5B. These observations demonstrate that our mathematical model is able to describe a theoretical network of validated interactions between miRNAs and TFs with systems-level properties mediating direct dosage compensation of MYC and STAT3, with a possible impact on STAT5A and STAT5B. If this model is correct, the inhibition of some miRNAs negatively controlling the expression levels of the target genes could potentially block gene dosage compensation.

### A minimal model of MYC and STAT3 dosage compensation leads to the identification of putative targets against aneuploid cancer

We next asked which are the minimal components of the network to identify the exact regulatory loops mediating gene dosage compensation and to design strategies to interfere with this mechanism in silico. First, we sought to identify the essential species involved in the feedback or feed-forward loops controlling gene dosage compensation. We performed parameter scans changing the copy number of the compensated genes (MYC and STAT3) and observed the behavior of the concentrations of the miRNAs and TFs of the model. For MYC copy number scan, the MYC expression showed a compensated increase, observed also for several TFs and miRNAs (figure 3A left), including those identified by metabolic control analysis (miR-19a, miR-20a, miR-17, figure 3E) but others, such as STAT3 and miR21, decreased with increasing MYC copy numbers (Figure 3A left). For STAT3, increasing gene copy numbers led to a compensated increase of STAT3, several miRNAs including miR21, miR17, miR221, miR19a and miR20a and other TFs but also a decrease in STAT5A, ETS1 and PPARG (Figure 3A right).

**Figure 3.**
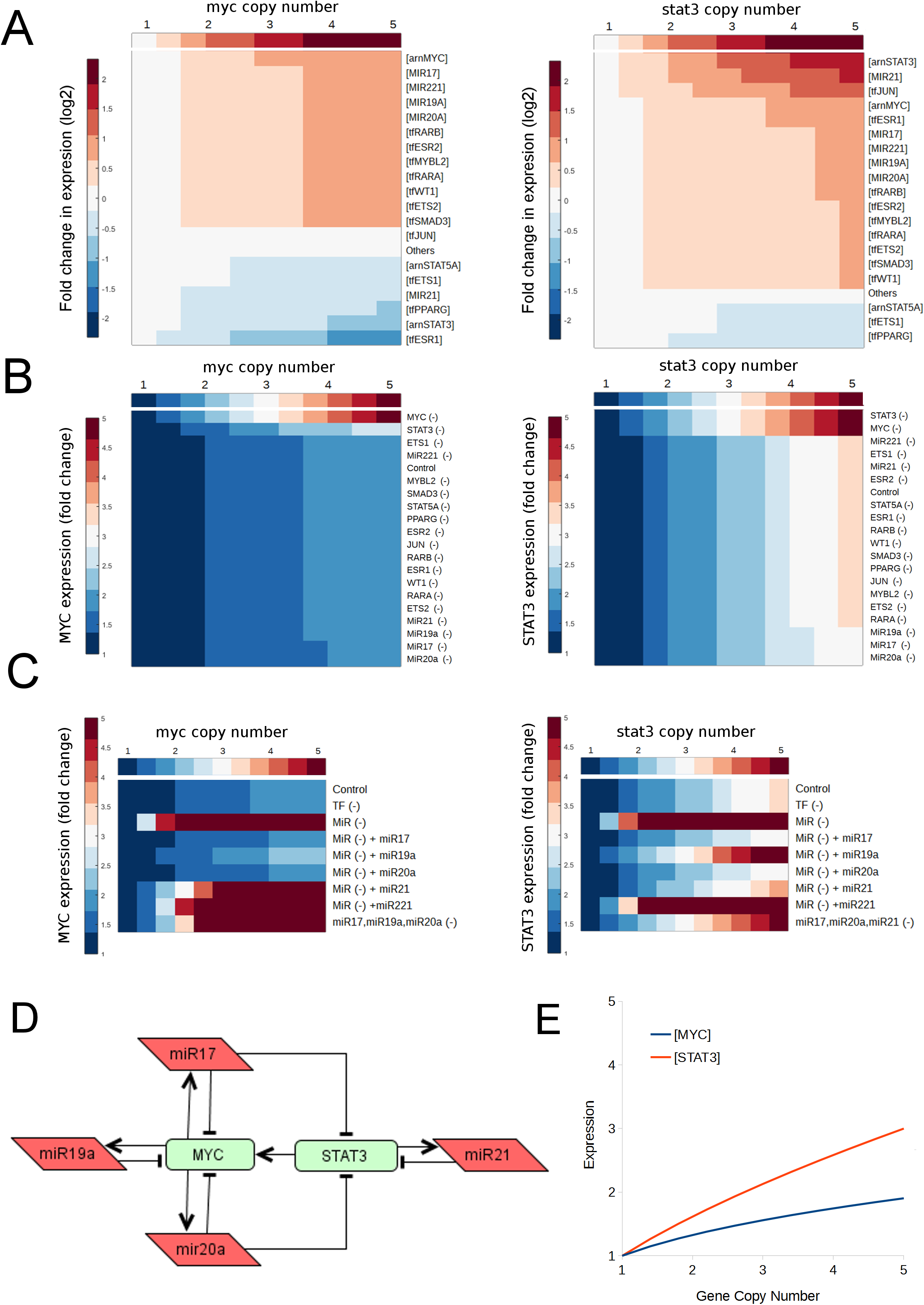
Biology-inspired experiments with the mathematical model led to the identification of a minimal model of gene dosage compensation for MYC and STAT3. A. A parameter scan on MYC and STAT3 copy numbers identifies which molecules increase or decrease together with varying copy numbers of those two genes. B. The single inhibition of those varying molecules has no effect on the gene dosage compensation of MYC and STAT3. C. The inhibition of groups of molecules and the restoration of single miRNAs indicate a redundant mechanism for the gene dosage compensation of MYC and STAT3. D. A minimal model of gene dosage compensation for MYC and STAT3 is hypothesized based on the results of these experiments. E. The fitting and parameter scan demonstrates that this minimal model recapitulates the dosage compensation of MYC and STAT3.

To assess the role in dosage compensation of these miRNAs and TFs responding the changes in MYC and STAT3 copy numbers, we performed single *in silico* inhibitions of the corresponding nodes by setting their interaction (inhibition/activation) parameters to 0. The inhibitions of single species altered the steady state levels of expression of MYC and STAT3 for the basal conditions (copy number of 1) but this effect is not shown due to the normalization of the expression, to better appreciate the effect on gene dosage compensation (figure 3B). The parameter scans under these conditions showed that MYC and STAT3 compensations are completely abolished when their respective interaction parameters are set to 0, confirming that their TF function is essential for the network to sense the changes in copy number (MYC(-) and STAT3(-) conditions, figure 3B). Also, MYC compensation is slightly altered when STAT3 is inhibited (STAT3(-)) and STAT3 compensation is completely abolished when MYC is inhibited (MYC(-), suggesting an interplay in the mechanisms of gene dosage compensation of these two genes. In contrast, the single inhibition for all remaining TFs and miRNAs had little or no effect on MYC and STAT3 dosage compensation, suggesting that this mechanism could be redundantly mediated by more than one miRNA or TF.

Therefore, we perform another in silico experiment changing the copy number of MYC and STAT3 while all miRNAs or TFs were inhibited, except by MYC and STAT3 themselves (Figure 4C). The results showed that the inhibition of all the other TFs (TF (-)) had no effect on the dosage compensation of MYC and STAT3. However, the inhibition of all miRNAs (MiR(-)) completely abolished the dosage compensation for both genes and led to a huge increase in their expression. We then proceeded to restore some miRNAs individually back into the MiR inhibited model, for example (MiR(-) + miR17) to simulate the effect of the active miR17 alone. Surprisingly, more than one miRNA was able to restore dosage compensation for both MYC and STAT3, confirming thereby that this mechanism is redundant. Indeed, miR-17, miR-19a and miR20a were individually able to compensate MYC dosage and miR17, miR20a and miR21 were able restore STAT3 dosage compensation (Figure 3C). Additionally, to confirm the role of these redundant miRNAs on gene dosage compensation in silico, we performed their triple inhibition in the complete model. Indeed, the triple inhibition (miR17, miR19a and miR20a (-)) was able to block MYC dosage compensation (figure 3C left), whereas (miR17, miR20a, miR21 (-)) abolished STAT3 dosage compensation (Figure 3C right), confirming that MYC and STAT3 dosage compensation is mediated by a redundant mechanism.

**Figure 4.**
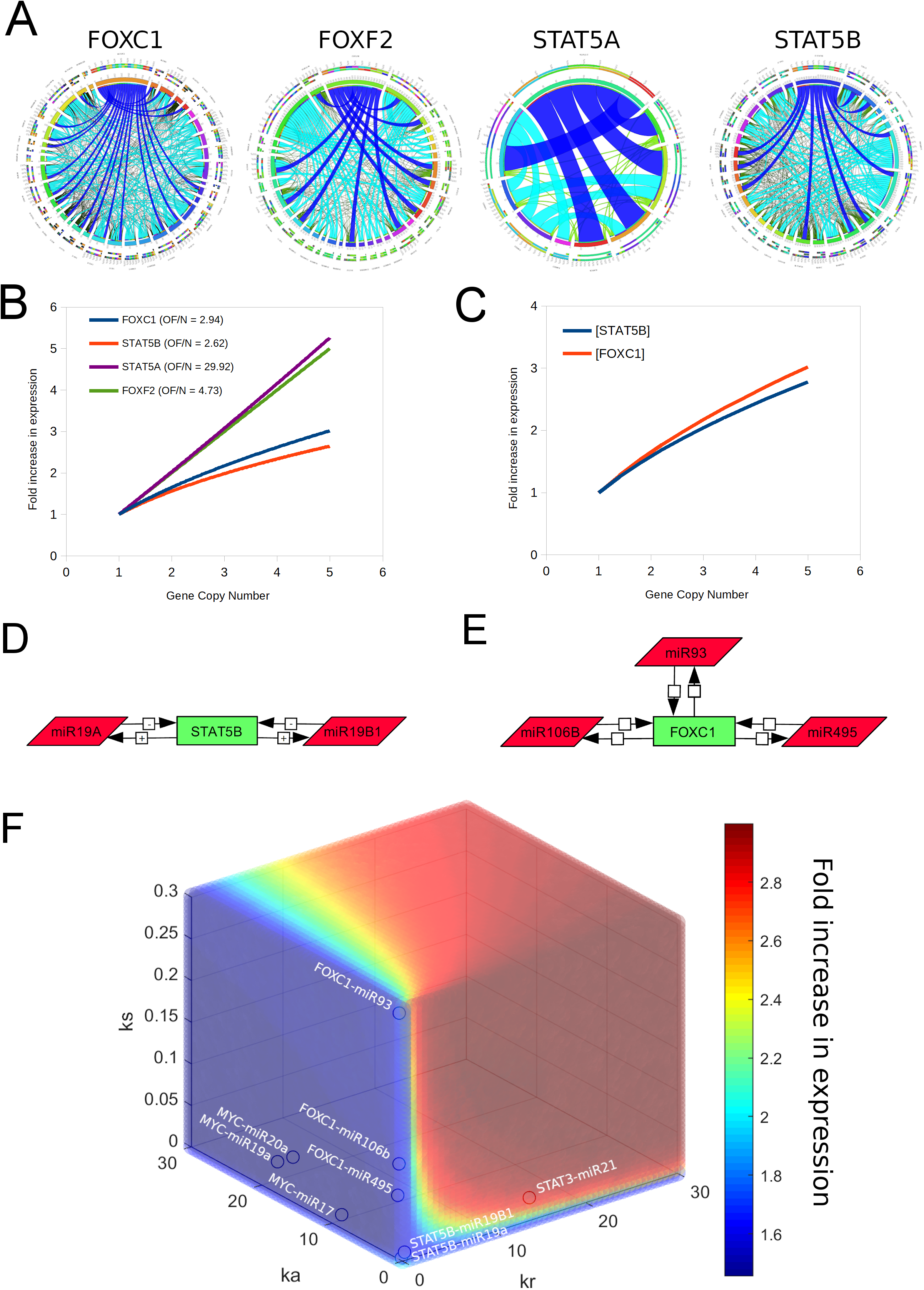
The gene dosage compensation depends on the kinetic parameters of the interactions of basic regulatory network motifs. A. Networks of direct interactions (putative and validated) of other candidate target genes with TFs and miRNAs. B. The fitted models show dosage compensation for STAT5B and FOXC1. C. The minimal models of STAT5B and FOXC1 recapitulate dosage compensation. D. The minimal model of STAT5B includes two redundant feedback loops with miR19A and miR19B1. E. The minimal model of FOXC1 dosage compensation includes three redundant feedback loops with miR106B, miR93 and miR495 F. A three dimensional landscape of gene dosage compensation dependent on the values of ks (synthesis rate of the miRNA), ka (activation parameter of the TF) and kr (repression parameter of the miRNA) demonstrates that the gene dosage compensation mediated by some basic regulatory network motifs depends on the values of these kinetic parameters.

Based on these results of redundant compensation and the interplay between MYC and STAT3, we proposed a minimal network topology of gene dosage compensation for MYC and STAT3 (Figure 3D). After reconstructing the interactions from the original network, the minimal network states that MYC is compensated by 3 redundant negative feedback loops formed with miR17, miR19a and miR20a. The compensation of STAT3 is mediated by 1 feedback loop with miR21 and 2 feed-forward loops: (STAT3-MYC-miR17-STAT3) and (STAT3-MYC-miR20a-STAT3). From this minimal network topology, we constructed a minimal mathematical model of gene dosage compensation for MYC and STAT3. After fitting this minimal model to the experimental data and repeating the parameter scan experiments for gene copy numbers of MYC and STAT3, we reconstituted the same dosage compensation mechanism observed in the original large model of interactions (Figure 3E). These results indicate that we identified *in silico* a minimal model of gene dosage compensation for MYC and STAT3 mediated by their interactions with miR17, miR19a, miR20a and miR21.

### The kinetic parameters of the TF-miRNA interactions determine the capability of a putative network motif for gene dosage compensation

Since we were able to identify a minimal model of gene-dosage compensation for MYC and STAT3 with their interactions with 4 miRNAs, we next asked whether we could identify common motifs of gene dosage-compensation for other TFs from our list of candidates. Due to the high complexity of the previous models, we could employ only experimentally-validated interactions to construct the networks, which probably precluded the identification of some motifs of gene dosage compensation for other less studied candidate genes. Indeed, the MYC-STAT3 model is data-driven but its identification was dependent on the available information of miRNA-TF-gene interactions: 3 negative feedback loops compensate MYC, whereas 1 negative feedback loop and 2 feed-forward loops compensate STAT3. These results suggest that the identification of those types of basic network motifs could lead to the identification of gene dosage compensation.

To continue with this rationale, we constructed the networks of all direct interactions (putative and validated) of 4 single genes with other TFs and miRNAs: FOXC1, FOXF2, STAT5A and STAT5B. Next, we filtered those arcs to include only negative feedback loops with miRNAs and feed-forward loops of 2 types: Gene-TF-miRNA-Gene and Gene-miRNA-TF-Gene. The resulting network topologies include 148 arcs for FOXC1, 114 arcs for FOXF2, 20 arcs for STAT5A and 97 arcs for STAT5B (figure 4A). These networks were subsequently converted into ODE models and the corresponding NCI60 data was extracted to proceed with the parameter estimation task in COPASI first using EP (Evolutionary Programming) and then the Hooke-Jeeves optimization algorithms to refine the fitting. The models were fitted until no parameter boundary alert was present and the objective functions were low for each of them (Figure 4B, OF/N objective function divided by the number of fitted values). From the resulting fitted models, the model of STAT5B and FOXC1 showed dosage compensation (Figure 4B). After performing similar experiments to those presented in figure 4, we determined that the main regulators of STAT5B dosage compensation are miR19A and miR19B1 (Figure 4D). For FOXC1, three miRNAs formed negative feed-back loops to redundantly compensate its gene dosage (Figure 4E). Indeed, after constructing and fitting these minimal models of STAT5B and FOXC1 dosage compensation, we were able to reconstitute the behavior of dosage compensation for both genes (Figure 4C).

We next asked what is particular about those few feedback loops accounting for dosage-compensation, while many others do not. We perform a triple parameter scan using the minimal model of MYC-STAT3. Since the compensation in that model is redundant, we inhibited miR19A and miR20a so that we can evaluate only the behavior of the negative feedback loop between MYC and miR17. Next, we monitored the concentration of MYC for two conditions: a MYC copy number of 1 and a MYC copy number of 3. Under these two conditions we scanned the effect of different values of ks (synthesis rate of the miRNA), ka (activation parameter of the TF) and kr (repression parameter of the miRNA). After calculating the ratio of those 2 conditions, we obtained a three dimensional landscape of gene dosage compensation with a color-coded heatmap of the fold increase in MYC expression upon a 3 fold increase in copy number (Figure 4F). The blue color corresponds to a region of dosage-compensation whereas the red region represents the linear increase of expression as a function of copy number. Indeed, when we located the positional values of the parameters corresponding to the compensating motifs, they all appeared in the blue region of compensation (Figure 4F) except by the interactions between STAT3 and miR21. These results confirmed that the simple identification of network motifs such as feedback or feed-forward loops is not enough to identify systems-level properties such as gene-dosage compensation. This behavior is indeed dependent on the data-driven identification of the right parameter values accounting for the emergence of this property of gene dosage compensation.

### The therapeutic potential of targeting gene dosage compensation against aneuploid cancer

Finally, we aimed to assess the therapeutic potential of the manipulation of the phenomenon of gene-dosage concentration to target aneuploid cancer. First, we look for the copy number values and the corresponding expression of the 3 genes identified hereby as compensated but in a larger collection of cancer cell lines. Therefore, we employed the data of CCLE comprising 937 cell lines (Ghandi et al., 2019). These cell lines include 586 cell lines with MYC amplification (63%), 409 cell lines with STAT3 amplification (44%), 407 cell lines with STAT5B amplification (43%) and 371 cell lines with FOXC1 amplification (40%) (Figure 5A). The trendlines suggest a behavior of gene dosage compensation for almost all the cell lines with copy number amplification, suggesting that most aneuploid cancer cells could be targeted by blocking gene dosage compensation.

**Figure 5.**
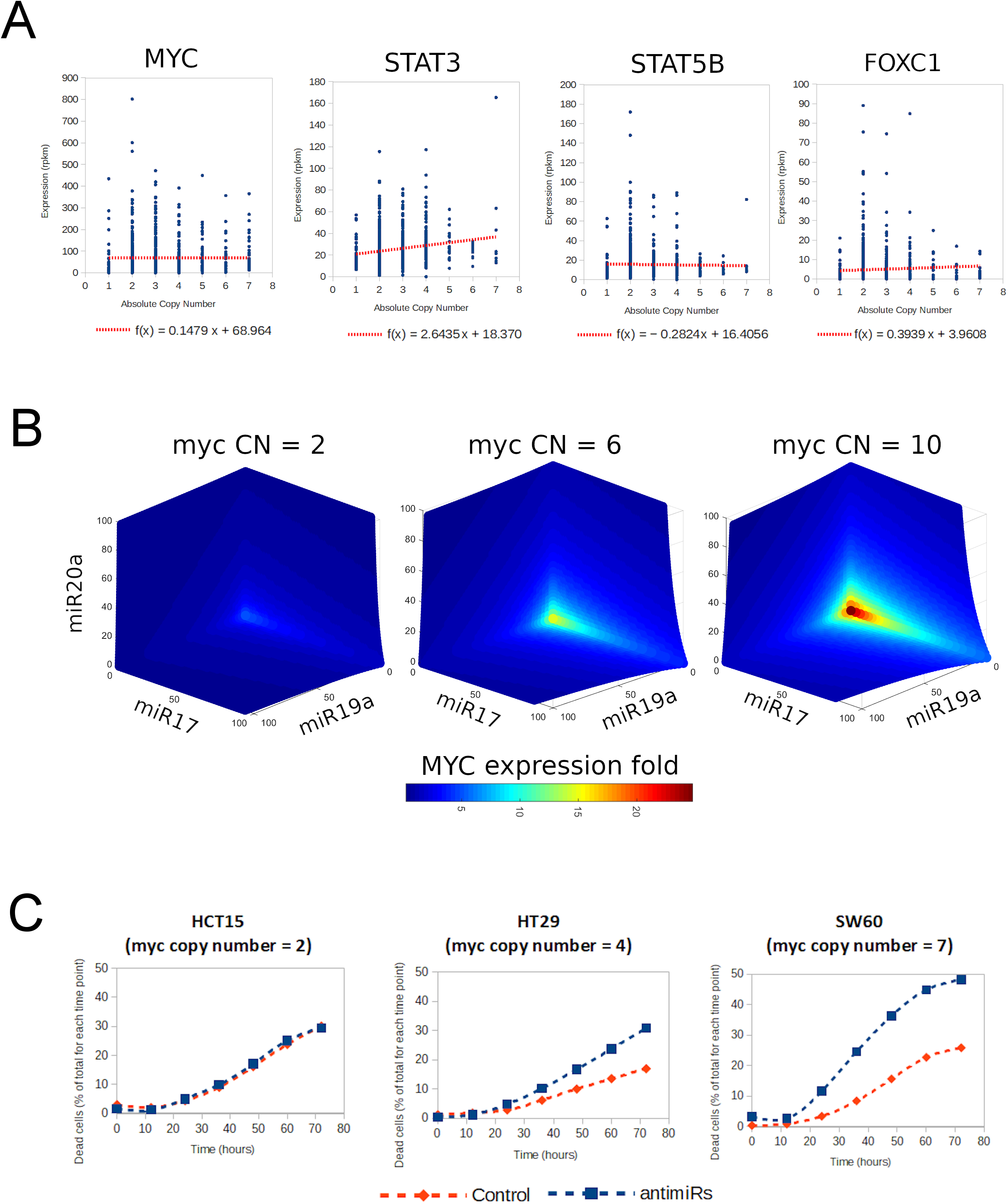
Therapeutic possibilities of targeting gene dosage compensation against aneuploid cancer. A. CCLE data shows copy number amplification in confirms 40-60% of cases, which confirms a behavior of gene dosage compensation for MYC, STAT3, STAT5B and FOXC1 (trendlines with sublinear slopes). B. The in silico simulations for the dependence of MYC concentration on the amounts of the 3 compensating miRNAs suggest that MYC dosage compensation in cancer with MYC amplification is more sensitive to the inhibition of those miRNAs. C. The experimental inhibition of those 3 miRNAs in colon cancer cells with 3 different copy numbers of MYC suggest a higher sensitivity to the inhibition of gene dosage compensation for more aneuploid cancers.

A second issue about the therapeutic potential of blocking gene dosage compensation is the specificity. Here, we focused on the dosage compensation of MYC. To block the redundant compensation of MYC, 3 miRNAs have to be blocked: miR17, miR19a and miR20a. How could this potentially impact diploid cells? In order to assess that, we performed a triple parameter scan on the ks (miRNA synthesis parameters) of the 3 miRNAs and plotted the MYC concentration values in function of the concentrations of those 3 miRNAs. The scan was repeated under 3 conditions of varying MYC copy numbers (2, 6 and 10) and the effect on the MYC concentration was observed (Figure 5B). Although there are some slight alterations under almost a complete depletion of the 3 miRNAs for the diploid condition (CN = 2), the increase in MYC copy number is much higher with increasing copy numbers of MYC, and less miRNA depletion is required. This result indicates that the perturbation of the mechanism of dosage compensation has a promising therapeutic range that increases with the extent of dosage amplification.

In order to confirm this hypothesis experimentally, we chose 3 colon cancer cell lines of the NCI60 panel with varying copy numbers of MYC, the HCT15 (2), the HT29 (4) and the SW60 (7). We chose a concentration of 3 anti-miRs combined to treat the cells and compared that with a scrambled RNA as control and monitored cell death over time until 70 hours. We observed no difference in the cell death induction by the antimiRs and the control RNA for the HCT15 cell line, whereas the difference between those two conditions increased with the copy number of MYC (figure 5C). These results indicates that cells with higher copy numbers of MYC are more sensitive to the blockade of gene dosage compensation by the perturbation of miR17, miR19a and miR20a. Altogether, our data-driven results report the transcriptional gene dosage compensation of MYC, STAT3 and STAT5B mediated by an emergent property of feedback and feed-forward loops. The interactions forming this network motifs have a specific set of kinetic parameters enabling compensation. This mechanism can be blocked by depleting key miRNAs, affecting in a larger extent those cells with higher copy numbers of those compensated genes, highlighting the potential of these kind of interventions to target aneuploid cancer.

## Discussion

Currently, the topic of miRNA regulation is gaining much attention in the field of molecular and cell biology. Indeed, the identification of a miRNA controlling cancer robustness has a huge therapeutic potential since miRNAs are becoming more attractive targets for therapy, as shown for the first time by miRNA-122 against hepatitis C infection and hepatic cancer (Lindow & Kauppinen, 2012). Nowadays, at least 7 miRNA mimics or inhibitors are tested in clinical trials thanks to the development of chemical modifications to increase their stability, to improve targeting to disease sites or the transport by several options of delivery systems (Rupaimoole & Slack, 2017). However, it is very hard to identify single miRNA-target interactions with relevant biological function. Classical approaches start with the identification of dysregulated miRNAs related to disease and extensive molecular and cellular biology work is required to validate the gene targets related to the phenotype of interest, which is inefficient because each miRNA can alter the expression of hundreds of genes by only 1.5 to 4 fold (Vera et al., 2013) and it is the cooperative effect of miRNA networks that makes them robust regulators (Herranz & Cohen, 2010; Matsuo, Eno, Im, Rosenshein, & Sood, 2010). Therefore, the identification of critical miRNAs likely to affect the response of those networks requires the analysis of a mathematical model of those interactions by means of sensitivity analysis combined with predictive simulations to suggest key biochemical processes to become potential therapeutic targets (Lai et al., 2013; Vera et al., 2013).

We developed a computational platform to construct large scale models of miRNAs and TF interactions with a reverse approach, starting with a list of target genes of interest. These genes could be determined by differential expression analysis or customized by the researchers. Afterwards, a network topology is constructed with all the reported interactions of those target genes with miRNAs and TFs. However, the network topology is not enough to identify the most robust targets to control the phenotype of interest. Therefore, our approach goes beyond identifying a network and establishes a full dynamical system based on ODEs, which is calibrated with existing experimental data. Finally, the sensitivity analysis and the in silico experiments enabled us to identify the most robust target nodes to regulate the phenotype of interest, to determine the mechanism behind that phenotype and the interventions on those target nodes.

In the present work we studied the mechanism of gene dosage compensation in aneuploid tumoral cells with the aid of models constructed by the biocomputational platform. First, we identified a list of candidate genes under transcriptional gene dosage compensation across the cancer genomes of the NCI-60 panel. Using these input genes we determined a miRNA/TF interaction network that connects all candidate genes but its structural analysis revealed that only those with transcription factor function can be subjected to gene dosage compensation. After model simplification, the targets with TF function were used to construct a mathematical model of “sensor loops” with all their reported and experimentally validated interactions with TFs and miRNAs. Within this model, an emergent property of gene dosage compensation was identified for MYC and STAT3 and a minimal model of MYC and STAT3 was extracted, determining that dosage compensation is mediated by feedback and feed-forward loops with 4 miRNAs. The kinetic parameters of these TF-miRNA interactions determine the capability of these network motifs to achieve gene dosage compensation. Searching for similar motifs among the putative interactions reported for other candidate target genes, we could suggest a putative mechanism for the dosage compensation of STAT5B and FOXC1 as well. Our results indicate that the miRNAs responsible for dosage compensation, particularly MYC, are putative targets against aneuploid cancer since this gene (and other candidates) are compensated in the larger CCLE data set. Finally, the in silico and in vitro inhibition of those 3 miRNAs indicates a higher sensitivity to perturbation in cells with MYC amplification pointing towards the therapeutic potential of targeting gene dosage compensation against aneuploid cancer.

Indeed, aneuploidy is lethal for normal cells and entire organisms with many negative effects given by the alteration of the gene dosage leading to unbalanced load of many proteins, some of them critical altering energetic requirements and protein homeostasis. However, not all of those dosage-altered genes are critical nor require compensation to maintain cell viability. Our data on the NCI60 panel shows that only a handful of genes have low tolerance to variation in their expression despite a high variation in their copy numbers (Figure 1). Indeed, others also reported that the messenger RNA (mRNA) levels generally correlate well with an increased DNA copy number (gene dosage) in aneuploid cancers (Stingele et al., 2012). We observed that behavior both at the protein and transcript level for our candidates pointing to a putative dosage compensation at the transcriptional level. However, the employed protein data set of the NCI60 is limited, probably precluding the identification of candidates compensated at the protein level only. In fact, gene dosage compensation was mostly reported at the protein level and not at the transcriptional level, whereby the stoichiometry of protein complexes in aneuploid cells determines that excess subunits are either degraded or aggregated to achieve dosage compensation (Brennan et al., 2019). Another work, using the genetic tug-of-war technique showed that approximately 10% of the genome shows gene dosage compensation at the protein level (Ishikawa et al., 2017), although this approach was not designed to distinguished compensation at the transcriptional level. To our knowledge, the only previous evidence of gene dosage compensation at the transcriptional level in cancer cells was provided for some few genes by a report of the insertion of an additional chromosome 5 (Stingele et al., 2012). The only report of dosage-compensation at the transcriptional level was done in wild yeast isolates compared to isogenic euploid strains, since laboratory strains are very sensitive to karyotypic changes (Hose et al., 2015).

The dosage compensation by the regulation of gene transcription might be an upstream and more effective mechanism, as it maintains the stoichiometry and preserves the energy required for transcription, translation, and eventual degradation of the extra proteins. Our approach was therefore intended to identify compensated candidates at the transcriptional level. To our knowledge, most reports try to identify potential therapeutic targets based on differential expression data between tumoral and normal tissues. Those differences arise as a consequence of genetic instability and as such, could be highly heterogeneous among cancer types. In contrast, our work is the first to explore stability of gene expression in cancer, represented by a handful of critical genes which could mediate cell survival despite cancer genomic instability. Indeed, aneuploidy is a hallmark of most advanced cancer cells that must have developed mechanisms to minimize its negative effects. As mentioned in the introduction, some conserved patterns in the karyotypic configurations suggest the presence of a stable mechanism, explained by the existence of a group of critical genes which function has to be maintained constant to ensure survival. We suggest that these essential genes establish a central core of stability that is selected or maintained within the evolutionary process of the unstable cancer genome. We also propose that this central core of stability is provided by a mechanism of gene dosage compensation of those critical genes, known to have very small tolerance to variation. While the expression of dosage-compensated genes is controlled in proportion to gene copy, many genes escape dosage compensation in response to aneuploidy and can contribute significantly to phenotypic variation in cancer.

The behavior of those candidate genes under compensation is therefore identified by our criteria of low tolerance in variation of their expression regardless of a high variation in their copy numbers. A similar criteria of a CNV-buffering score was already proposed for aneuploid yeast strains as the dosage-compensated genes had higher variation in copy number but a constraint gene expression (Hose et al., 2015). Indeed, our list of candidates includes many cancer-important genes, specially 7 transcription factors highly connected to many other genes (Table 1). Some of our candidate genes are clustered in common chromosomal locations, which is expected since copy number variants occur in large fragments of chromatin. However, the fact that they seemed to be all compensated suggests that the compensation of 1 gene within one of those clusters could be sufficient to co-regulate the expression of other connected genes, but this needs further work and larger data sets to be confirmed.

**Table 1.** Candidate genes under dosage compensation.

| Gen ID | Location | Description | Transcription factor | Forward loops | Type of regulation |  |
| --- | --- | --- | --- | --- | --- | --- |
|  |  |  |  |  | RNA | Protein |
| MYC | 8q24.21 | v-myc avian myelocytomatosis viral oncogene homolog | yes | 461 | yes | no data |
| ZNF217 | 20q13.2 | zinc finger protein 217 | yes | 451 | yes | yes |
| MTSS1 | 8p22 | metastasis suppressor 1 | no | 102 | yes | no data |
| SEMA3D | 7q21.11 | sema domain, immunoglobulin domain (Ig), short basic domain, secreted, (semaphorin) 3D | no | 40 | yes | yes |
| TRIM37 | 17q23.2 | tripartite motif containing 37 | no | 54 | yes | no data |
| STAT3 | 17q21.31 | signal transducer and activator of transcription 3 (acute-phase response factor) | yes | 577 | yes | yes |
| KCNH4 | 17q21.2 | potassium voltage-gated channel, subfamily H (eag-related), member 4 | no | 9 | yes | no data |
| RAB5C | 17q21.2 | RAB5C, member RAS oncogene family | no | 99 | yes | yes |
| FOXC1 | 6p25 | forkhead box C1 | yes | 41 | yes | no data |
| BIRC2 | 11q22 | baculoviral IAP repeat containing 2 | no | 2 | yes | yes |
| CUL5 | 11q22.3 | cullin 5 | no | 179 | yes | yes |
| ATM | 11q22-q23 | ATM serine/threonine kinase | no | 2 | yes | yes |
| NPAT | 11q22-q23 | nuclear protein, ataxia-telangiectasia locus | no | 76 | yes | yes |
| MMP12 | 11q22.3 | matrix metalloproteinase 12 (macrophage elastase) | no | 2 | yes | no data |
| DCUN1D5 | 11q22.3 | DCN1, defective in cullin neddylation 1, domain containing 5 | no | 31 | yes | yes |
| PGR | 11q22-q23 | progesterone receptor | yes | 1 | yes | yes |
| PDCD10 | 3q26.1 | programmed cell death 10 | no | 139 | yes | yes |
| ATP1B2 | 17p13.1 | ATPase, Na<sup>+</sup>/K<sup>+</sup> transporting, beta 2 polypeptide | no | 17 | yes | no data |
| ANKFY1 | 17p13.3 | ankyrin repeat and FYVE domain containing 1 | no | 147 | yes | yes |

We presumed that the behavior of the genes establishing that stability core would be very homogeneous among cells and cancer types in terms of compensation. Although the NCI60 panel has few cell lines (59) to argue on that, we could confirm a stable behavior of dosage compensation for most of the cases of amplified MYC, STAT3, stat5b and foxc1 for the CCLE data set that includes next generation data of more than one thousand cancer cell lines of many different cancer types (Ghandi et al., 2019). The stability in this behavior suggested a tightly regulation at the gene expression level. Indeed, miRNA networks are robust regulators of gene expression upon environmental changes (Herranz & Cohen, 2010) and they show adaptation to gene dosage through the formation of regulatory circuits with transcription factors (Bleris et al., 2011). Since the miRNA-TF networks are very complex, the gene dosage compensation could arise as an emergent property of the system meaning that a differential expression analysis of gene targets or miRNAs would be insufficient to identify the underlying mechanism of gene dosage compensation.

Therefore, we seeked to identify gene dosage compensation mechanisms mediated by the emerging properties of complex miRNA-TF regulatory networks. With the aid of the biocomputational platform we studied the gene dosage compensation mechanism in the NCI60 panel of cancer cell lines. This includes a collection of 7 types of cancer which are fully characterized at genomic and transcriptomic level and these data was an input for us to develop a biocomputational platform to model large-scale miRNA-TF networks. We first constructed a large-scale network of interactions including new sources for both putative and experimentally-validated interactions. Using this network, we constructed and fitted an ODE mathematical model, which was not able to reproduce any behavior of dosage compensation. An important characteristic in biological networks like the miRNA-TF networks is its capability to adapt to fluctuating concentrations of its biomolecular components. Coherent and incoherent feed forward loops (FFL) and feedback loops (FBL) help the network to adapt from such fluctuations (Carignano, Mukherjee, Singh, & Seelig, 2019; Osella, Bosia, Corá, & Caselle, 2011). Indeed, the feedback regulations were previously proposed to explain the gene dosage compensation in aneuploid yeast strains (Hose et al., 2015). Despite of the initial step-backs, the turning point toward the modeling of the gene dosage compensation was the insight of the sensor loop: the group of interactions starting in a transcription factor and returning after a defined number of interactions with other species including miRNAs and other TFs. These includes feedback and feed-forward loops but also more complex interactions. The sensor loop itself also helps the network adapt to fluctuations of the genes by repressing its expression. Once we added sensor loops in the models, we easily modeled this mechanism and validated the hypothesis for the genes MYC and STAT3 (Figure 2).

The gene MYC (MYC proto-oncogene, bHLH transcription factor), aka c-MYC, is a well known and studied transcription factor. It activates genes involved in proliferation, cell growth, cell differentiation and apoptosis. It is estimated that MYC regulates 15% of the genes. It is also an oncogene, usually overexpressed in many kinds of cancer (Dang, 1999) and it is active in 70% of human cancer but it is also related to apoptosis (Prendergast, 1999). The desregulation of MYC may lead cancer but may also lead to a cell suicide (Nilsson & Cleveland, 2003), and its reported to have a dual function from oncogene to tumor suppresor in leukemia (Uribesalgo, Benitah, & Di Croce, 2012).

STAT3, signal transducer and activator of transcription 3, is a TF that relay external signals to the nucleus where the signal regulates genes involved in differentiation, proliferation, apoptosis, angiogenesis, metastasis and immune system (Johnston & Grandis, 2011; Levy & Lee, 2002). In cancer cells this gene suppresses detection mechanism of the immune system, promotes angiogenesis, invasion and metastasis, deregulates growth (Yue & Turkson, 2009). STAT3 is an oncogene but recent information indicates that it may also function as a tumor suppressor (Avalle, Camporeale, Camperi, & Poli, 2017).

A deeper analysis of the mathematical model for MYC and STAT3 enabled us to understand the underlying mechanism of gene dosage compensation by the reduction of model complexity into a minimal model recapitulating the same behavior of dosage compensation for both genes (Figure 3). First, a sensitivity analysis in COPASI enabled us to identify the main TFs and miRNAs regulating the concentrations (but not the compensation) of MYC and STAT3. Second, a parameter scan varying the copy number of MYC and STAT3 enabled us to confirm which species increase or decrease together with increasing copy numbers for both genes. Afterwards, the same experiment was repeated along with the single inhibition of those species but the compensation behavior was unaltered. This immediately indicated that the mechanism of compensation is redundant, prompting us to inhibit groups of species instead. The inhibition of all miRNAs indeed abolished gene dosage compensation but it was restored by the single reactivation of 3 miRNAs for each gene. Using the wild type model we could confirm that the only way to abolish gene dosage compensation was to simultaneously inhibit those 3 corresponding miRNAs for each gene. After reconstructing the interactions from the original network, the minimal network states that MYC is compensated by 3 redundant negative feedback loops formed with miR17, miR19a and miR20a. The compensation of STAT3 is mediated by 1 feedback loop with miR21 and 2 feed-forward loops: (STAT3-MYC-miR17-STAT3) and (STAT3-MYC-miR20a-STAT3)(Figure 3). The miR17, miR19a and miR20a are actually all co-regulated as the miRNA cluster miR-17-92. This is considered an oncogenic cluster and is actually called OncomiR-1 since it is overexpressed in several types of cancer (for review see (Fuziwara & Kimura, 2015)). Several reports showed that these miRNAs form important network motifs with MYC in B-cell lymphoma (Mihailovich et al., 2015), with E2F/MYC (Y. Li, Li, Zhang, & Chen, 2011) and with STAT3 in retinoblastoma (Jo et al., 2014). This biology-inspired in silico experiments enabled us to propose a minimal model of gene dosage compensation for MYC and STAT3 out of experimentally-validated interactions. Due to the redundancy in gene dosage compensation, it would be unfeasible to identify such mechanism by single or even double inhibitions using a functional genomics approach without the guidance of a systems biology approach.

The biocomputational platform developed was an accelerator of discovery for this work. By constructing models in minutes and automating several processes the platform shortened the time required to complete the construction of mathematical models. The compilation of around 65,000 experimentally validated regulatory interactions had a huge impact on the project. The amount of interactions allowed the platform to build interesting models with enough interactions. And by not using putative interactions, the models were cleaner and smaller.

Nevertheless, our model has been simplified in several ways. We included only experimentally validated interactions in the model presenting gene dosage compensation. From the 21 candidate genes, our current model offers an explanation for the gene dosage compensation of MYC and STAT3 only. This is clearly a limitation due to the small amount of cases of the NCI60 panel representing only 59 cases of aberrant genomes. Further work with larger data sets such as the CCLE and TCGA could lead to the identification of further candidate genes and the availability of more data to obtain more accurate models of gene dosage compensation. In addition, we faced a limitation due to paramount complexity of the mathematical models, which prevent us from using putative (not experimentally validated) interactions to reduce model complexity. Despite of these limitations, and based on the experience with the minimal model for MYC and STAT3, we built single gene models for other four compensated candidates including both putative and experimentally validated interactions. Hereby, we filtered for only those arcs establishing FBL or FFL obtaining smaller models suitable for a faster parameter estimation. Among those, we could provide a putative explanation (out of putative interactions) for the gene dosage compensation of STAT5B and FOXC1 and reconstruct a minimal model for their corresponding underlying mechanisms of compensation (Figure 4). Interestingly, MYC, STAT3, STAT5B and FOXC1 present a behavior similar to dosage compensation on the CCLE data (Figure 5A). When plotting the kinetic parameters describing the main interactions of the compensating network motifs for all those genes within a three dimensional landscape of gene dosage compensation, we observed that they are all located in the compensation region. Therefore, we critically established that the network topology alone is not enough, as with different parameter values the model would lose gene dosage compensation, strengthening the need for full dynamic models.

The broad compensation of those target genes in CCLE and the further analysis of the current model of gene dosage compensation of MYC and STAT3 or even other genes could reveal novel specific targets against cancer. Thus, we suggest that cancer has a robust Achilles-Heel due to an increased sensitivity to perturbations in these circuits, which is not necessarily reflected as differences in miRNA expression levels but at systems-level properties. The identification of control points blocking the dosage compensation could lead to the over-expression of these two genes and others under their influence in a context of fragility for the cancer cell. These strategy is promising inasmuch the overexpression of these important transcription factors seem to be more sensitive to blockade of gene dosage compensation when their copy number are more amplified as expected in silico (Figure 5B) but also suggested by the higher cytotoxicity of colon cancer cells with higher MYC copy numbers upon inhibition of the three miRNAs proposed here to mediate its dosage compensation. Indeed, dysregulated MYC triggers rapid apoptosis (Nilsson & Cleveland, 2003) and STAT3 may function as a tumor suppressor (Avalle et al., 2017). An important prediction of the study of gene dosage compensation in aneuploid yeast was indeed that the phenomenon of dosage compensation occurs at genes that are most toxic when overexpressed (Hose et al., 2015).

In conclusion, the present work led to the construction of a complex mathematical model to study gene dosage compensation and formulated model-driven hypothesis for the identification of novel targets against aneuploid cancer. In addition, the computational platform built with this project has other potential applications to understand miRNA-mediated gene regulation and perform simulations of the systems-level effects of perturbations in miRNA networks related to disease. Furthermore, since current techniques such as Next Generation Sequencing allows the rapid acquisition of genomic data for cancer patients, this platform can be easily adapted to generate personalized computer models to identify the optimal targets according to each cancer configuration. Future work is required to confirm the effect of gene dosage compensation on patient survival and to identify other compensation cores to direct personalized precision therapies against cancer.

Altogether, the current results could contribute to the identification of that stability core of essential genes, which manipulation of specific nodes has the potential to become a novel approach to specifically target aneuploid cancer cells.

## Materials and Methods

### A. Data sources

Data was gathered from several sources. The primary sources were from experiments on the NCI60 panel: gene copy number (Bussey et al., 2006), RNA gene expression (Shankavaram et al., 2007) and protein expression (Gholami et al., 2013). MicroRNA related data was downloaded from Mirtarbase (Hsu et al., 2014) and MiRBase (Kozomara et al., 2019; Kozomara & Griffiths-Jones, 2014). For background knowledge on gene regulation we relied on several sources: Transmir (Wang, Lu, Qiu, & Cui, 2010), Pazar (Portales-Casamar et al., 2009), TRED (Transcriptional Regulatory Element Database) (Jiang, Xuan, Zhao, & Zhang, 2007), CircuitsDB (Friard, Re, Taverna, De Bortoli, & Corá, 2010).

### B. Gene classification using Gaussian Mixture Models

In order to classify genes according to their behavior, we developed a computer algorithm based on the Gaussian Mixture Model functions in MATLAB. An increasing number of components (ki) of the GMM model is added sequentially and the GMM training is performed for several iterations searching for the best fit to the experimental data. The resulting GMM is used to classify the cells of the original data set. A MANOVA test is applied to the resulting clusters to evaluate the statistical significance of adding another component to the GMM. This is done until the new component adds no further significance.

### C. Construction of the Regulatory Network

We are interested in the regulatory network of miRNAs and TFs in the nearness of and directly affecting the genes selected by the GMM. In order to build this network, we relied heavily on the interaction database gathered previously. A regulatory interaction in this database is an arc which starting point is the regulator and the ending point is the regulatee. For many interactions, the database tells whether the regulation is an activation or repression. When the database does not provide this information, if the regulator of the interaction is a TF we assume that the regulation is an activation, and if the regulator is a miRNA we assume that it is a repression.

We use a directed graph to represent this network. We first identify the nodes of this graph which are composed by all the genes selected by the GMM as well as all the direct regulators and regulated nodes of these genes. To identify the regulators, we search in the interaction database for arcs ending in one of the selected genes. The direct regulators are all the starting nodes of each of the arcs found. Similarly, we search for arcs starting in one of the selected genes and the end node of the arc is a directly regulated gene. Hence, we have successfully identified all the nodes of the graph. Each node is a selected gene, a miRNA or TF, all of them biomolecular species relevant in the gene dosage compensation phenomenon. We then build the list of arcs of the graph. From the list of nodes or species identified, for each pair of species, if there is an arc in the interaction database linking these two species, we add this arc in the list of arcs of the graph. The lists of species and arcs identified becomes the graph representing the regulatory network affecting the selected genes of interest.

In order to examine the target/miRNA/TF network for the presence of regulatory motifs with systems-level properties, we searched for positive and negative feedback loops (between miRNAs and TFs), coherent feed-forward loops and incoherent feed-forward loops.

### D. Ordinary differential equation modeling of miRNA-TF networks

We now construct a system of ODEs from the graph representation of the regulation network. Since the gene dosage compensation phenomenon is gene expression related, we are interested in constructing a metabolic model from the graph. For each species of the graph, we write an equation computing the concentration or mRNA expression of this specie. The list of all these equations defines a system of ODEs modeling the metabolic behavior of the regulation network. We have three different types of equations, one for each type of species: candidate gene, TF and miRNA. The schematic representations of these equations are shown in (Figure 2B).

For a candidate gene, the mRNA expression equation has a synthesis part and a degradation part. The expression of each gene equals to the synthesis part subtracted by the degradation part.

The synthesis equation of the candidate gene is the product of three factors: a) the gene copy number, b) the synthesis rate of the gene, a parameter to be fitted by the Parameter Estimation process, and c) the total regulatory effect exercised by TFs on the candidate gene which in the graph is represented by all the incoming arcs of the candidate gene where the arc’s regulator is a TF. The total regulatory effect factor is usually >1 since most TFs are activators but it could be <1 when the regulating TFs are mainly repressors.

The degradation equation of a candidate gene is also the product of three factors: a) the mRNA concentration of the gene, b) the degradation rate of the gene, another parameter to be estimated, and c) the total regulatory effect of miRNAs on the gene. Because the degradation equation is a product and the total repression effect is one of the factors, the value of the repression effect is usually >1 since miRNAs are repressors, thus exercising an accelerating effect on the degradation.

For modeling the total regulatory effect, we define the following parameters, also to be estimated later against experimental data. For a TF we define an activation rate that represents the rate at which the TF activates o intensifies the expression of the TF’s regulatees. Likewise we define a repression rate representing the rate at which the TF suppresses or silences its regulatees when the TF has a repressing role. For a miRNA we also define a repression rate since miRNA can only repress its regulatees. The single regulatory effect of one TF or one miRNA over all of its regulatees is the product of the concentration of the TF or miRNA multiply by the corresponding rate. For instance, the concentration of MYC multiplied by MYC’s activation rate or the concentration of mir19 by mir19’s repression rate. Given that the concentration of a regulator, the activation rate and the repression rate are all zero or positive, the single effect of one regulator is also zero or positive. Following this definition, we have only one repression rate for a miRNA, therefore a given miRNA will exercise the same rate of repression over each of its regulatees.

We model the total repression effect of miRNAs in the degradation equation of a candidate gene as the sum of 1 plus the single effect of each of the miRNAs repressing that gene. This value is always greater or equal than 1. When there are no miRNA repressing the gene, this factor would be 1. When there is at least one miRNA repressing the gene, this factor would be greater than one as long as the concentration of the miRNA is positive. These values model the expected function of the total repression effect in the degradation equation for the presence of repressing miRNAs would accelerate the degradation of the gene.

We define the total regulation effect of TFs in the synthesis equation of a candidate gene as a ratio between the total activation effect of the TFs divided by the total repression effect of the TFs. The total activation effect is the sum of 1 plus the single effect of each of the TFs activating the gene. Likewise, the total repression effect is the sum of 1 plus the single effect of each TFs repressing the gene. When there are no TFs regulating the gene, this ratio is 1. When there are TFs activating or repressing the gene, if the total activation effect is greater than the total repression effect then the ratio is greater than 1, hence accelerating the synthesis of the gene. If there are more repression effect than activation effect, the ratio is less than 1 slowing down the synthesis. If both effects are equal, the ratio is 1 exercising no effect on the synthesis. These ratio’s values model the expected behavior of the interplay between the activating and repressing TFs in the synthesis equation of a gene.

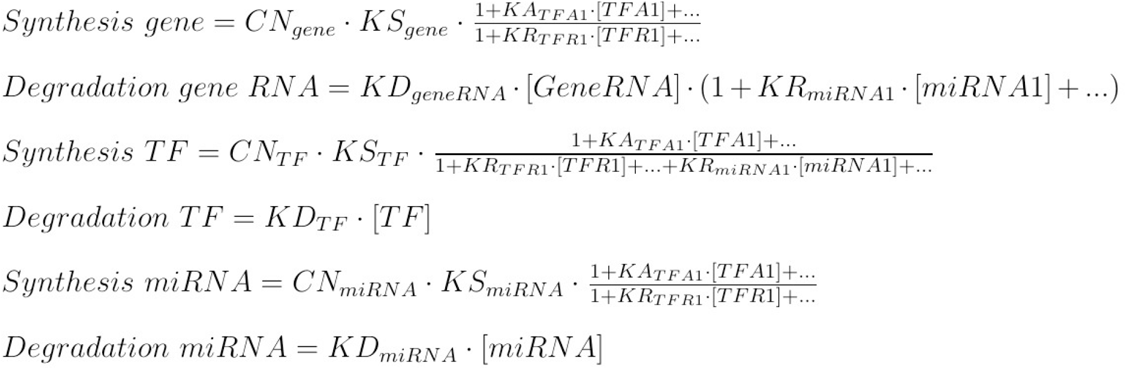

The mRNA expression equations for a TF or miRNA are slightly different from the equation of a candidate gene. These two equations also have a synthesis part and a degradation part but instead of having the repressing effect exercising an accelerator effect in the degradation part, they have the repressing effect in the synthesis part applying a decelerating effect in the synthesis part.

The degradation equation for a TF or miRNA is the product of only two factors: a) the mRNA concentration of the TF or miRNA, and b) the degradation rate of TF or miRNA. The synthesis equation for a TF or miRNA is the product of three factors: a) the copy number of the TF or miRNA, b) the synthesis rate of the TF or miRNA, and c) the total regulatory effect exercised by others TFs and miRNAs on this TF or miRNA which in the graph is represented by all the incoming arcs of this TF or miRNA where the arc’s regulator is a TF or miRNA.

While the total activation effect depends only on activating TFs, now the total repressing effect involves not only repressing TFs but also repressing miRNAs. Whether there are more total activating effect or more total repressing effect, the total regulatory effect would be >1 or <1, or exactly 1 if both effects are the same. Although this equation applies to TFs and miRNAs, in practice the synthesis equation for miRNA do not involve repressing miRNA effects since miRNAs do not repress other miRNAs.

From the three types of equations defined, we detect three different types of elements: copy numbers, rates and concentrations. There is a copy number parameter for each species of the graph. There are four different types of rates o parameters to be fitted against experimental data: synthesis, degradation, activation and repression rates. A candidate gene has only a synthesis and degradation rate. A TF has all four types of rates. And a miRNA has all but an activation rate since miRNAs only repress. Finally, there is also one concentration for each species of the graph.

The concentrations of the regulators that are part of the total regulation effect factor of the equation play an important role in the modeling of the regulatory effects. For an activator TF, as its concentration increases, so increases the activation effect and the concentration of the TF’s regulated gene. In the same way, as the concentration of a miRNA increases, so increases the repression effect causing a decline in the concentration of the miRNA’s regulated gene, either by slowing down its synthesis or by accelerating its degradation. The concentrations also play an important role in the modeling of common network motifs like loops like feedback loops, feedforward loops or other types of loops. In a loop like A->B->C->A, the concentration of A is part of the regulatory factor of B’s equation, the concentration of B is part of C’s equation, and C’s concentration is part of A’s equation. The interplay of these three equations models the dynamics of the loop in the graph.

Having defined the equations of the species and identified the different elements of these equations, translating the graph of the regulatory network into an SMBL model (Hucka et al., 2003) is straightforward. An SBML model follows the XML standard (W3C/XML, 2008) and have several sections for the different elements of the SBML model. The two most relevant sections are the parameters section and the equation sections. In the parameter section we add the copy number and the rates of all the species of the graph. In the reaction section we add one reaction for each species of the graph. Since we have all the information needed for the SBML model, constructing the model is just a matter of filling correctly the different sections of the model.

### E. Model parameter estimation

The SBML model is imported into COPASI, an application for analyzing biochemical networks like the miRNA-TF regulatory network. The platform also add 59 sets of experimental data to this model, one set for each cell line of NCI60. We then run a *Parameter Estimation* task in COPASI. The task adjusts the parameters of the model to fit the RNA expression of each species of the model as close as possible to the corresponding RNA expression of 59 experimental data-sets.

To assess that the RNA expression of the model is *close enough* to the RNA expression of the experimental data: a) we build a model for each cell line by using the fitted model and changing the gene copy number and initial concentration values to the values of the corresponding cell line; b) we run a *Time Course* task for each model and gather the RNA expression for each species in each of the cell lines; c) we use a dependent *t*-test for paired samples with α=0.05 to compare the model’s RNA expression of the 60 cell lines of a species with the corresponding experimental RNA expression. The pairing is on the cell lines. Species with significant differences between their experimental and model’s RNA expression are discarded from the model and a new Parameter Estimation task is run for the modified model until the difference between the experimental and the simulated data is not significant (T-student, p > 0.05). The presumption is that the experimental data do not explain the discarded species and they therefore do not belong to the model.

### G. Assessment of gene dosage compensation behavior

To evaluate if a gene is compensated in the fitted model, we run a *Parameter Scan* in COPASI, varying the copy number of the gene from 1 to 5 and computing the respective RNA expression. A non-compensated gene has a near linear fold increase in RNA expression level as its corresponding copy number increases linearly. We consider a gene compensated if the increase in RNA expression follows a sublinear increase as the copy number increases linearly.

### H. Trendline CCLE MYC, STAT3, STAT5B and FOXC1 genes calculation

In order to calculate the absolute copy number for MYC, STAT3, STAT5B and FOXC1, we use the calculated absolute copy number (Carter et al., 2012) analysis data of the CCLE (Ghandi et al., 2019) and the human Genome Reference Consortium build 38 (Schneider et al., 2017) as inputs of a program written in python that takes the gene location from the genome reference to search its copy number value in the copy number analysis data. After that, for each gene we use the copy number and the CCLE expression data together with the least squares method to calculate the trendline, finally we plotted the expression against the copy number and the calculated trendline.

## Acknowledgements

We thank the initial contribution of Dr. Michael Kesling for his advises on the project formulation and the data analysis. This project was funded by the grant FEES-CONARE (Costa Rica) and Vicerrectoría de Investigación, University of Costa Rica and Universidad Estatal a Distancia (UNED).

## Author contributions

Acón, ManSai^1,2^. Oviedo, Guillermo^1,2^. Baéz, Edwin^1,2^. Vásquez-Vargas, Gloriana^1^, Guevara-Coto, José^3^. Segura-Castillo, Andrés^6^. Siles-Canales, Francisco^4,5^. Quirós-Barrantes, Steve^1,5^. Mendes, Pedro^7^. Mora-Rodríguez, Rodrigo^1,2,5^.

### Conceptualization

A,M. QB, S. MR,R. Data curation: A,M. O,G. B,E. Formal Analysis: A,M. O,G. B,E. SC,F. M,P. Funding acquisition: MR, R. Investigation: A,M. SC,A. SC,F, MR,R. Methodology: A,M. VV, G. SC, A. Project administration: QB, S. MR, R. Resources: SC, F. QB,S. SC, A. M,P. MR,R. Software: A,M. O,G. B,E. GC, J. SC, A. Supervision: QB,S. MR, R. Validation: VV, G. M, P. Visualization: A,M. SC, F. MR,R. Writing – original draft: A,M. MR,R. Writing – review & editing: M,P.

Expanded View Figure 1. DNA, RNA and protein levels for all candidate genes.

Expanded View Figure 2. Putative miRNA-transcription factor interaction network linking all candidate target genes. A. Correlation coefficients of copy number variations of the candidate genes and miRNA/transcription factor expression across the NCI60 panel. B. Correlation coefficients of copy number variations of the candidate genes and miRNA/transcription factor expression across the NCI60 panel, only for those putative interactions present on the network in C. C. Network of putative interactions among target genes, miRNAs and transcription factors (red indicates repression and blue activation) including different types of regulatory loops formed by target genes, miRNAs and TFs.

## Reference List

Avalle, L., Camporeale, A., Camperi, A., & Poli, V. (2017). STAT3 in cancer: A double edged sword. Cytokine. https://doi.org/10.1016/j.cyto.2017.03.018

Birchler, J. a, & Veitia, R. a. (2012). Gene balance hypothesis: connecting issues of dosage sensitivity across biological disciplines. Proceedings of the National Academy of Sciences of the United States of America, 109(37), 14746–14753. https://doi.org/10.1073/pnas.1207726109

Bleris, L., Xie, Z., Glass, D., Adadey, A., Sontag, E., & Benenson, Y. (2011). Synthetic incoherent feedforward circuits show adaptation to the amount of their genetic template. Molecular Systems Biology, 7(519), 1–12. https://doi.org/10.1038/msb.2011.49

Blower, P. E., Verducci, J. S., Lin, S., Zhou, J., Chung, J.-H., Dai, Z., … Sadee, W. (2007). MicroRNA expression profiles for the NCI-60 cancer cell panel. Molecular Cancer Therapeutics, 6(5), 1483–1491. https://doi.org/10.1158/1535-7163.MCT-07-0009

Bond, U., Neal, C., Donnelly, D., & James, T. C. (2004). Aneuploidy and copy number breakpoints in the genome of lager yeasts mapped by microarray hybridisation. Current Genetics, 45(6), 360–370. https://doi.org/10.1007/s00294-004-0504-x

Brennan, C. M., Vaites, L. P., Wells, J. N., Santaguida, S., Paulo, J. A., Storchova, Z., … Amon, A. (2019). Protein aggregation mediates stoichiometry of protein complexes in aneuploid cells. Genes & Development, 33(15–16), 1031–1047. https://doi.org/10.1101/gad.327494.119

Bussey, K. J., Chin, K., Lababidi, S., Reimers, M., Reinhold, W. C., Kuo, W.-L., … Weinstein, J. N. (2006). Integrating data on DNA copy number with gene expression levels and drug sensitivities in the NCI-60 cell line panel. Molecular Cancer Therapeutics, 5(4), 853–867. https://doi.org/10.1158/1535-7163.MCT-05-0155

Carignano, A., Mukherjee, S., Singh, A., & Seelig, G. (2019). Extrinsic Noise Suppression in Micro RNA Mediated Incoherent Feedforward Loops. Proceedings of the IEEE Conference on Decision and Control. https://doi.org/10.1109/CDC.2018.8619371

Carter, S. L., Cibulskis, K., Helman, E., McKenna, A., Shen, H., Zack, T., … Getz, G. (2012). Absolute quantification of somatic DNA alterations in human cancer. Nature Biotechnology. https://doi.org/10.1038/nbt.2203

Dang, C. V. (1999). c-Myc Target Genes Involved in Cell Growth, Apoptosis, and Metabolism. Molecular and Cellular Biology. https://doi.org/10.1128/mcb.19.1.1

Devlin, R. H., Holm, D. G., & Grigliatti, T. a. (1982). Autosomal dosage compensation Drosophila melanogaster strains trisomic for the left arm of chromosome 2. Proceedings of the National Academy of Sciences of the United States of America, 79(4), 1200–1204. Retrieved from http://www.pubmedcentral.nih.gov/articlerender.fcgi?artid=345929&tool=pmcentrez&rendertype=abstract

Donnelly, N., & Storchová, Z. (2014). Dynamic karyotype, dynamic proteome: buffering the effects of aneuploidy. Biochimica et Biophysica Acta, 1843(2), 473–481. https://doi.org/10.1016/j.bbamcr.2013.11.017

Duesberg, P., Rausch, C., Rasnick, D., & Hehlmann, R. (1998). Genetic instability of cancer cells is proportional to their degree of aneuploidy. Proceedings of the National Academy of Sciences of the United States of America, 95(23), 13692–13697. https://doi.org/10.1073/pnas.95.23.13692

Fabarius, A., Li, R., Yerganian, G., Hehlmann, R., & Duesberg, P. (2008). Specific clones of spontaneously evolving karyotypes generate individuality of cancers. Cancer Genetics and Cytogenetics, 180(2), 89–99. https://doi.org/10.1016/j.cancergencyto.2007.10.006

Fabarius, A., Willer, A., Yerganian, G., Hehlmann, R., & Duesberg, P. (2002). Specific aneusomies in Chinese hamster cells at different stages of neoplastic transformation, initiated by nitrosomethylurea. Proceedings of the National Academy of Sciences of the United States of America, 99(10), 6778–6783. https://doi.org/10.1073/pnas.251670699

Fabian, M. R., Sonenberg, N., & Filipowicz, W. (2010). Regulation of mRNA translation and stability by microRNAs. Annual Review of Biochemistry, 79, 351–379. https://doi.org/10.1146/annurev-biochem-060308-103103

Friard, O., Re, A., Taverna, D., De Bortoli, M., & Corá, D. (2010). CircuitsDB: a database of mixed microRNA/transcription factor feed-forward regulatory circuits in human and mouse. BMC Bioinformatics, 11, 435. https://doi.org/10.1186/1471-2105-11-435

Fuziwara, C. S., & Kimura, E. T. (2015). Insights into regulation of the miR-17-92 cluster of miRNAs in cancer. Frontiers in Medicine, 2(SEP), 13–17. https://doi.org/10.3389/fmed.2015.00064

Ghandi, M., Huang, F. W., Jané-Valbuena, J., Kryukov, G. V., Lo, C. C., McDonald, E. R., … Sellers, W. R. (2019). Next-generation characterization of the Cancer Cell Line Encyclopedia. Nature. https://doi.org/10.1038/s41586-019-1186-3

Gholami, A. M., Hahne, H., Wu, Z., Auer, F., Meng, C., Wilhelm, M., & Kuster, B. (2013). Global proteome analysis of the NCI-60 cell line panel. Cell Reports, 4(3), 609–620. https://doi.org/10.1016/j.celrep.2013.07.018

Gmeiner, W. H., Reinhold, W. C., & Pommier, Y. (2010). Genome-wide mRNA and microRNA profiling of the NCI 60 cell-line screen and comparison of FdUMP[10] with fluorouracil, floxuridine, and topoisomerase 1 poisons. Molecular Cancer Therapeutics, 9(12), 3105–3114. https://doi.org/10.1158/1535-7163.MCT-10-0674

Hanahan, D., & Weinberg, R. A. (2011). Hallmarks of cancer: The next generation. Cell, 144(5), 646–674. https://doi.org/10.1016/j.cell.2011.02.013

Hanna, J., Hossain, G. S., & Kocerha, J. (2019). The potential for microRNA therapeutics and clinical research. Frontiers in Genetics, 10(MAY). https://doi.org/10.3389/fgene.2019.00478

Herranz, H., & Cohen, S. M. (2010). MicroRNAs and gene regulatory networks: managing the impact of noise in biological systems. Genes & Development, 24(13), 1339–1344. https://doi.org/10.1101/gad.1937010

Hooke, R., & Jeeves, T. A. (1961). “Direct Search” Solution of Numerical and Statistical Problems. Journal of the ACM (JACM). https://doi.org/10.1145/321062.321069

Hose, J., Yong, C. M., Sardi, M., Wang, Z., Newton, M. A., & Gasch, A. P. (2015). Dosage compensation can buffer copynumber variation in wild yeast. ELife. https://doi.org/10.7554/eLife.05462

Hsu, S. Da, Tseng, Y. T., Shrestha, S., Lin, Y. L., Khaleel, A., Chou, C. H., … Huang, H. Da. (2014). MiRTarBase update 2014: An information resource for experimentally validated miRNA-target interactions. Nucleic Acids Research, 42(D1). https://doi.org/10.1093/nar/gkt1266

Hucka, M., Finney, A., Sauro, H. M., Bolouri, H., Doyle, J. C., Kitano, H., … Wang, J. (2003). The systems biology markup language (SBML): A medium for representation and exchange of biochemical network models. Bioinformatics. https://doi.org/10.1093/bioinformatics/btg015

Ishikawa, K., Makanae, K., Iwasaki, S., Ingolia, N. T., & Moriya, H. (2017). Post-Translational Dosage Compensation Buffers Genetic Perturbations to Stoichiometry of Protein Complexes. PLoS Genetics, 13(1). https://doi.org/10.1371/journal.pgen.1006554

Jiang, C., Xuan, Z., Zhao, F., & Zhang, M. Q. (2007). TRED: A transcriptional regulatory element database, new entries and other development. Nucleic Acids Research, 35(SUPPL. 1). https://doi.org/10.1093/nar/gkl1041

Jo, D. H., Kim, J. H., Cho, C. S., Cho, Y. L., Jun, H. O., Yu, Y. S., … Kim, J. H. (2014). STAT3 inhibition suppresses proliferation of retinoblastomathrough down-regulation of positive feedback loop of STAT3/miR-17-92 clusters. Oncotarget. https://doi.org/10.18632/oncotarget.2546

Johnston, P. A., & Grandis, J. R. (2011). STAT3 signaling: Anticancer strategies and challenges. Molecular Interventions. https://doi.org/10.1124/mi.11.1.4

Kim, D., Sung, Y. M., Park, J., Kim, S., Kim, J., Park, J., … Baek, D. (2016). General rules for functional microRNA targeting. Nature Genetics, 48(12), 1517–1526. https://doi.org/10.1038/ng.3694

Kitano, H. (2004). Cancer as a robust system: implications for anticancer therapy. Nature Reviews. Cancer, 4(3), 227–235. https://doi.org/10.1038/nrc1300

Kozomara, A., Birgaoanu, M., & Griffiths-Jones, S. (2019). MiRBase: From microRNA sequences to function. Nucleic Acids Research, 47(D1), D155–D162. https://doi.org/10.1093/nar/gky1141

Kozomara, A., & Griffiths-Jones, S. (2014). MiRBase: Annotating high confidence microRNAs using deep sequencing data. Nucleic Acids Research, 42(D1). https://doi.org/10.1093/nar/gkt1181

Kvitek, D. J., Will, J. L., & Gasch, A. P. (2008). Variations in stress sensitivity and genomic expression in diverse S. cerevisiae isolates. PLoS Genetics, 4(10), 31–35. https://doi.org/10.1371/journal.pgen.1000223

Lai, X., Bhattacharya, A., Schmitz, U., Kunz, M., Vera, J., & Wolkenhauer, O. (2013). A systems’ biology approach to study microrna-mediated gene regulatory networks. BioMed Research International, 2013(Ii). https://doi.org/10.1155/2013/703849

Levy, D. E., & Lee, C. K. (2002). What does Stat3 do? Journal of Clinical Investigation. https://doi.org/10.1172/JCI0215650

Li, L., McCormack, A. a, Nicholson, J. M., Fabarius, A., Hehlmann, R., Sachs, R. K., & Duesberg, P. H. (2009). Cancer-causing karyotypes: chromosomal equilibria between destabilizing aneuploidy and stabilizing selection for oncogenic function. Cancer Genetics and Cytogenetics, 188(1), 1–25. https://doi.org/10.1016/j.cancergencyto.2008.08.016

Li, Y., Li, Y., Zhang, H., & Chen, Y. (2011). MicroRNA-mediated positive feedback loop and optimized bistable switch in a cancer network Involving miR-17-92. PloS One, 6(10), e26302. https://doi.org/10.1371/journal.pone.0026302

Lindow, M., & Kauppinen, S. (2012). Discovering the first microRNA-targeted drug. The Journal of Cell Biology, 199(3), 407–412. https://doi.org/10.1083/jcb.201208082

Matsuo, K., Eno, M. L., Im, D. D., Rosenshein, N. B., & Sood, A. K. (2010). Gynecologic Oncology Clinical relevance of extent of extreme drug resistance in epithelial ovarian carcinoma. Gynecologic Oncology, 116(1), 61–65. https://doi.org/10.1016/j.ygyno.2009.09.018

Mihailovich, M., Bremang, M., Spadotto, V., Musiani, D., Vitale, E., Varano, G., … Bonaldi, T. (2015). MiR-17-92 fine-tunes MYC expression and function to ensure optimal B cell lymphoma growth. Nature Communications. https://doi.org/10.1038/ncomms9725

Neph, S., Stergachis, A. B., Reynolds, A., Sandstrom, R., Borenstein, E., & Stamatoyannopoulos, J. A. (2012). Circuitry and dynamics of human transcription factor regulatory networks. Cell, 150(6), 1274–1286. https://doi.org/10.1016/j.cell.2012.04.040

Nilsson, J. A., & Cleveland, J. L. (2003). Myc pathways provoking cell suicide and cancer. Oncogene. https://doi.org/10.1038/sj.onc.1207261

Osella, M., Bosia, C., Corá, D., & Caselle, M. (2011). The role of incoherent microRNA-mediated feedforward loops in noise buffering. PLoS Computational Biology. https://doi.org/10.1371/journal.pcbi.1001101

Ozery-Flato, M., Linhart, C., Trakhtenbrot, L., Izraeli, S., & Shamir, R. (2011). Large-scale analysis of chromosomal aberrations in cancer karyotypes reveals two distinct paths to aneuploidy. Genome Biology, 12(6), R61. https://doi.org/10.1186/gb-2011-12-6-r61

Park, Y., & Kuroda, M. I. (2001). Epigenetic aspects of X-chromosome dosage compensation. Science, 293(5532), 1083–1085. https://doi.org/10.1126/science.1063073

Portales-Casamar, E., Arenillas, D., Lim, J., Swanson, M. I., Jiang, S., McCallum, A., … Wasserman, W. W. (2009). The PAZAR database of gene regulatory information coupled to the ORCA toolkit for the study of regulatory sequences. Nucleic Acids Research, 37(SUPPL. 1). https://doi.org/10.1093/nar/gkn783

Prendergast, G. C. (1999). Mechanisms of apoptosis by c-Myc. Oncogene. https://doi.org/10.1038/sj.onc.1202727

Ritchie, W., Rasko, J. E. J., & Flamant, S. (2013). MicroRNA target prediction and validation. Advances in Experimental Medicine and Biology, 774, 39–53. https://doi.org/10.1007/978-94-007-5590-1_3

Rupaimoole, R., & Slack, F. J. (2017). MicroRNA therapeutics: Towards a new era for the management of cancer and other diseases. Nature Reviews Drug Discovery, 16(3), 203–221. https://doi.org/10.1038/nrd.2016.246

Schneider, V. A., Graves-Lindsay, T., Howe, K., Bouk, N., Chen, H. C., Kitts, P. A., … Church, D. M. (2017). Evaluation of GRCh38 and de novo haploid genome assemblies demonstrates the enduring quality of the reference assembly. Genome Research. https://doi.org/10.1101/gr.213611.116

Shankavaram, U. T., Reinhold, W. C., Nishizuka, S., Major, S., Morita, D., Chary, K. K., … Weinstein, J. N. (2007). Transcript and protein expression profiles of the NCI-60 cancer cell panel: an integromic microarray study. Molecular Cancer Therapeutics, 6(3), 820–832. https://doi.org/10.1158/1535-7163.MCT-06-0650

Sheltzer, J. M., & Amon, A. (2011). The aneuploidy paradox: costs and benefits of an incorrect karyotype. Trends in Genetics, 27(11), 446–453. https://doi.org/10.1016/j.tig.2011.07.003

Shimoga, V., White, J. T., Li, Y., Sontag, E., & Bleris, L. (2014). Synthetic mammalian transgene negative autoregulation. Molecular Systems Biology, 9(1), 670–670. https://doi.org/10.1038/msb.2013.27

Solé, R. V, & Deisboeck, T. S. (2004). An error catastrophe in cancer? Journal of Theoretical Biology, 228(1), 47–54. https://doi.org/10.1016/j.jtbi.2003.08.018

Stingele, S., Stoehr, G., Peplowska, K., Cox, J., Mann, M., & Storchova, Z. (2012). Global analysis of genome, transcriptome and proteome reveals the response to aneuploidy in human cells. Molecular Systems Biology, 8(608), 608. https://doi.org/10.1038/msb.2012.40

Tong, Z., Cui, Q., Wang, J., & Zhou, Y. (2019). TransmiR v2.0: An updated transcription factor-microRNA regulation database. Nucleic Acids Research. https://doi.org/10.1093/nar/gky1023

Uribesalgo, I., Benitah, S. A., & Di Croce, L. (2012). From oncogene to tumor suppressor: The dual role of Myc in leukemia. Cell Cycle. https://doi.org/10.4161/cc.19883

Veitia, R. A., Bottani, S., & Birchler, J. A. (2008). Cellular reactions to gene dosage imbalance : genomic, transcriptomic and proteomic effects. (June). https://doi.org/10.1016/j.tig.2008.05.005

Vera, J., Lai, X., Schmitz, U., & Wolkenhauer, O. (2013). MicroRNA-regulated networks: the perfect storm for classical molecular biology, the ideal scenario for systems biology. Advances in Experimental Medicine and Biology, 774, 55–76. https://doi.org/10.1007/978-94-007-5590-1_4

Wang, J., Lu, M., Qiu, C., & Cui, Q. (2010). TransmiR: a transcription factor-microRNA regulation database. Nucleic Acids Research, 38(Database issue), D119–22. https://doi.org/10.1093/nar/gkp803

Yue, P., & Turkson, J. (2009). Targeting STAT3 in cancer: How successful are we? Expert Opinion on Investigational Drugs. https://doi.org/10.1517/13543780802565791

